# Acetazolamide modulates intracranial pressure directly by its action on the cerebrospinal fluid secretion apparatus

**DOI:** 10.1101/2022.01.11.475854

**Authors:** Dagne Barbuskaite, Eva K. Oernbo, Jonathan H. Wardman, Trine L. Toft-Bertelsen, Eller Conti, Søren N. Andreassen, Niklas J. Gerkau, Christine R. Rose, Nanna MacAulay

## Abstract

Elevated intracranial pressure (ICP) is observed in many neurological pathologies, e.g. hydrocephalus and stroke. This condition is routinely relieved with neurosurgical approaches, since effective and targeted pharmacological tools are still lacking. The carbonic anhydrase inhibitor, acetazolamide (AZE), may be employed to treat elevated ICP. However, its effectiveness is questioned, its location of action unresolved, and its tolerability low. Here, we employed *in vivo* and *ex vivo* approaches to reveal the efficacy and mode of action of AZE in the rat brain. The drug effectively reduced the ICP, irrespective of the mode of drug administration and level of anaesthesia. The effect occurred via a direct action on the choroid plexus and an associated decrease in cerebrospinal fluid secretion, and not indirectly via the systemic action of AZE on renal and vascular processes. Upon a single administration, the reduced ICP endured for approximately 10 h post-AZE delivery with no long-term changes of brain water content or choroidal transporter expression. However, a persistent reduction of ICP was secured with repeated AZE administrations throughout the day. Future specific targeting of choroidal carbonic anhydrases may limit the systemic side effects, and therefore enhance the treatment tolerability and effectiveness in select patient groups experiencing elevated ICP.

## Introduction

Elevated intracranial pressure (ICP) is observed in various brain pathologies. It causes spatial compression of the brain tissue and reduced cerebral perfusion, and if left untreated, leads to cerebral ischaemia and may be fatal (1). Treatment of elevated ICP relies heavily on surgical interventions, most commonly ventriculoperitoneal shunting in hydrocephalic patients, and craniectomy in severe cases of brain edema (2). Only a few pharmacological options are available for relief of elevated ICP, the most widely used being Diamox®, with the active ingredient acetazolamide (AZE) (3). AZE is an inhibitor of the carbonic anhydrases, the 15 isoforms of which are widely expressed in different cell types and tissues throughout the mammalian body (4). These enzymes catalyse the reversible conversion of CO_2_ to H_2_CO_3_, which is followed by its dissociation to hydrogen ion (H^+^) and bicarbonate (HCO_3_ ). AZE was launched in the early 1950s as a diuretic agent (5), and later employed for treatment of elevated ICP (6). However, although some clinical studies revealed AZE-mediated ICP decrease (7), others demonstrated modest, if any, effect of AZE treatment (8, 9). Due to its wide expression, AZE treatment associates with a range of systemic side effects, e.g. paraesthesia, dysgeusia, polyuria and fatigue (10). The uncertainty regarding AZE efficiency, combined with poor patient compliance, has raised questions regarding its usability in the clinic (11). Notwithstanding, due to lack of alternatives, AZE remains a first-line treatment for certain ICP pathologies, e.g. idiopathic intracranial hypertension (12), irrespective of the unresolved mode of action by which the drug may confer relief of elevated ICP. Studies on experimental animals have remained equally inconclusive as AZE administration did (13, 14) or did not (15) reduce the ICP.

In contrast to its unclear effects on ICP, there is solid preclinical data supporting AZE’s efficacy in lowering cerebrospinal fluid (CSF) secretion in humans (16, 17) as well as in different experimental animal models, i.e. sheep (18), dogs (19), rabbits (20–24), cats (25–28) and rats (29–31) (for review, see (32)). The majority of the CSF is secreted by the choroid plexus (33), which is an epithelial monolayer with a polarised expression of various ion transporters and channels (34, 35). Amongst these choroidal membrane transport mechanisms are several HCO_3_^-^ transporters, i.e. the sodium-driven chloride bicarbonate exchanger (NBCn2/NCBE) (36) and the anion exchanger 2 (AE2) (37) residing in the basolateral membrane, and the sodium bicarbonate cotransporter 2 (NBCe2) expressed in the luminal membrane (38, 39), all of which may be implicated in CSF secretion (40, 41). The activity of these transporters is determined by the availability of their substrate (HCO_3_ ). Inhibition of the choroidal carbonic anhydrases (42–46), and the ensuing reduction in [HCO_3_ ]_i_ in the choroid plexus tissue, therefore directly affects the transport rate and the associated CSF secretion. However, the rate of CSF secretion could also be modulated as a secondary effect of carbonic anhydrase inhibition elsewhere in the body. An example of such AZE-mediated modulation of physiological parameters occurs in the kidney, where carbonic anhydrase-mediated HCO_3_ conversion is required for the reabsorption of HCO_3_ (47). Inhibition of renal carbonic anhydrase might thus indirectly reduce the activity of the HCO_3_ transporters in choroid plexus by simply reducing blood HCO_3_ , thereby limiting access to their transported substrate. In addition, the AZE-mediated reduction in renal HCO_3_ reabsorption causes diuresis that, in turn, reduces the mean arterial blood pressure (MAP) (48, 49). Alteration in MAP is proposed to indirectly affect CSF secretion (50), and could thus lower the ICP in this manner. Furthermore, carbonic anhydrases in erythrocytes and the surrounding capillary endothelium supports efficient CO_2_ exchange between the tissue-blood-alveoli (51). Inhibition of such vascular carbonic anhydrases causes systemic elevation of the partial carbon dioxide pressure (pCO_2_) (52, 53) and ensuing hyperventilation, which may indirectly decrease the ICP (54). Uncontrolled breathing of anaesthetized animals that are not mechanically ventilated during the experimental procedure may thereby represent a confounding element to the former preclinical studies evaluating AZE-mediated effects on CSF secretion or ICP (55).

AZE is thus potentially a clinically useful pharmacological agent employed to reduce ICP in neurological conditions. However, it is unclear, if the well-established AZE-mediated reduction in CSF secretion does in fact lead to an associated reduction in ICP. Another key question is whether the reduced CSF secretion arises from AZE’s direct effect on choroidal carbonic anhydrases or occurs secondarily to AZE’s modulatory effect on other physiological processes such as blood pressure, kidney function, and/or blood gas content. Here, we demonstrate by complementary *ex vivo* and *in vivo* experimental approaches, conducted on both anesthetized and awake rats, that AZE lowers the ICP in healthy rats by its *direct action* on the choroidal carbonic anhydrases and subsequent reduction in CSF secretion. This new insight may guide future pharmacological treatment of elevated ICP targeted specifically to the choroid plexus.

## Methods

### Animals

Experiments were conducted in 9-10 week old male Sprague Dawley rats, housed in a temperature-controlled room with a 12h:12h light-dark cycle (6am to 6pm), and with free access to a standard rodent pellet diet and tap water. Animals were randomly allocated to each treatment group, and all experimental work was performed and reported in compliance with the ARRIVE guidelines (56).

### AZE and control solution formulation

For i.v. administration, AZE (A6011, Sigma-Aldrich) was dissolved in 5N NaOH to a 700 mg ml^-1^ stock solution, which was diluted in 0.9% NaCl to a working concentration of 20 mg ml^-1^, pH 8.4. The control solution (vehicle) was an equiosmolar 1.4% NaCl solution, pH 8.4. The solutions (5 ml kg^-1^ animal) were injected into either the femoral or the tail vein, resulting in a dose of 100 mg AZE kg^-1^ animal. For the intracerebroventricular (i.c.v.) delivery, AZE was dissolved in DMSO (1-2.5 M), and further diluted in either HCO_3_ -containing artificial CSF (aCSF; (in mM) 120 NaCl, 2.5 KCl, 2.5 CaCl_2_, 1.3 MgSO_4_, 1 NaH_2_PO_4_, 10 glucose, 25 NaHCO_3_, pH adjusted with 95% O_2_/5% CO_2_) or in HEPES-buffered artificial CSF (HEPES-aCSF; (in mM) 120 NaCl, 2.5 KCl, 2.5 CaCl_2_, 1.3 MgSO_4_, 1 NaH_2_PO_4_, 10 glucose, 17 Na-HEPES (4-(2-hydroxyethyl)-1-piperazineethanesulfonic acid), pH 7.4), when the solution could not be equilibrated with 95% O_2_/5% CO_2_. Depending on the experimental paradigm, the final working solutions were formulated to expose the choroid plexus tissue with a 200-500 µM AZE concentration, which should block carbonic anhydrases by 99.99% (57). Vehicle consisted of matching DMSO concentration (maximum 0.1%) with addition of mannitol when required to obtain equiosmolar solutions. For chronic oral (p.o.) administration, a suspension of AZE (in 0.9% NaCl, 50 mg ml^-1^) was delivered to the experimental rats in quantities (0.9-1.0 ml) to result in 100 mg kg^-1^ animal AZE once daily for 7 days (at 10am) or 3 times per day (at 7am, 2pm and 9pm) for 5 days by oral gavage. Control animals received saline via oral gavage.

### Anesthesia and physiological parameter monitoring

All non-survival surgeries were performed in rats anaesthetized with xylazine and ketamine (ScanVet, 10 mg kg^-1^ animal xylazine, 5min later 100 mg kg^-1^ animal ketamine, half dose of ketamine was re-dosed every 10-40min upon detection of foot reflex). Body temperature was maintained at 37°C by a homeothermic monitoring system (Harvard Apparatus). In experiments lasting for more than 30min, rats were tracheostomized and mechanically ventilated with the VentElite system (Harvard Apparatus), inhaling 0.9 l min^-1^ humidified air mixed with 0.1 l min^-1^ O_2_. The ventilation was adjusted according to exhaled end tidal CO_2_ (etCO_2_), measured with a capnograph (Type 340, Harvard Apparatus), to result in 5.0 ± 0.5 kPa blood pCO_2_ before administration of control or drug solutions. In the hyperventilation experiments, the ventilation used during the baseline period was increased by 50% (both respiratory rate and the tidal volume), and then maintained at these levels throughout the 2h experiment. The MAP was monitored through a heparinized saline-filled (15 IU heparin ml^-1^ in 0.9% NaCl) catheter inserted into the femoral artery, connected to a pressure transducer APT300, and transducer amplifier module TAM-A (Hugo Sachs Elektronik). The blood pressure signal was recorded at a 1 kHz sampling rate using BDAS Basic Data Acquisition Software (Hugo Sachs Elektronik). This catheter also served for blood sample collection required for blood gas determination with an ABL80 (Radiometer). All survival surgeries were performed under aseptic conditions on rats anesthetized with isoflurane (Attane vet, 1000 mg g^-1^ isoflurane, ScanVet), using 5% isoflurane (mixed with 1.8 l min^-1^ air / 0.2 l min^-1^ O_2_) in the anesthesia induction chamber, and 1-2.5% isoflurane to maintain anesthesia through a face mask throughout the surgery. The body temperature was maintained at 37°C by a homeothermic monitoring system (Harvard Apparatus).

### ICP recordings in anesthetized rats

Anesthetized and ventilated rats, placed in a stereotactic frame, had the skull exposed, and a 3.6 mm diameter cranial window drilled with care not to damage the dura. The epidural probe (PlasticsOne, C313G) was secured with dental resin cement (Panavia SA Cement, Kuraray Noritake Dental Inc.) and the ICP cannula was filled with HEPES-aCSF before connection to a pressure transducer APT300 and transducer amplifier module TAM-A (Hugo Sachs Elektronik). To ensure the presence of a continuous fluid column between the dura and the epidural probe, approximately 5 µl HEPES-aCSF was injected through the epidural probe. The ICP signal was recorded at a 1 kHz sampling rate using BDAS Basic Data Acquisition Software (Hugo Sachs Elektronik). Jugular compression was applied to confirm proper ICP recording. In the ICP recording during i.c.v. delivery of test solutions, a 0.5 mm burr hole was drilled contralateral to the ICP probe (1.3 mm posterior, 1.8 mm lateral to bregma), and a 4 mm brain infusion cannula (Brain infusion kit2, Alzet) placed into the lateral ventricle. Upon stabilization of the ICP signal, 37°C aCSF + 0.9% DMSO was infused (0.5 µl min^-1^) with a peristaltic pump for 25min prior to solution shift to either control solution (aCSF + mannitol + 0.9% DMSO) or AZE (aCSF + 18 mM AZE in 0.9% DMSO; expected ventricular concentration of 500 µM, equivalent to an accumulated 0.24 mg animal^- 1^ over the experimental timeline) for 120min. In the ICP experiments with i.v. delivery of test solutions, these were injected into the femoral vein through a heparinized saline-filled (15 U heparin ml^-1^ in 0.9% NaCl) catheter (100 mg kg^-1^ animal).

### Nephrectomy

Nephrectomy was performed on anesthetized rats by ligation (with 4-0 non-absorbable suture) of the renal arteries and veins after dorsal entry through the muscle layer. The incision sites were closed with metal wound clamps (Michel, 11×2mm) after the ligation.

### LI-COR live imaging

Anesthetized rats in a stereotaxic frame had their cranium and upper neck muscles exposed, and a burr hole drilled (same coordinates as i.c.v. cannula placement in ICP recordings) into which a Hamilton syringe (RN 0.40, G27, a20, Agntho’s) was inserted 4 mm into the lateral ventricle. The experiment was initiated with intraventricular injection (1.5 µl s^-1^) of 15 µl aCSF containing either vehicle (0.1% DMSO) or AZE (2 mM; expected ventricular concentration of 200 µM due to dilution in ∼150 µl native CSF). The procedure was repeated after 5min, but with inclusion of carboxylate dye (10 µM; IRDye 800CW, P/N 929-08972, LI-COR Biosciences). In experiments with i.v. delivery of AZE (100 mg kg^-1^ animal), these were injected into the tail vein through a catheter (24 G Neoflon, VWR) 25min prior to carboxylate dye injection. Image acquisition was initiated 1min after carboxylate injection and continued for 5min with 30s intervals using a Pearl Trilogy Small Animal Imaging System (LI-COR) (800 nm channel, 85 µm resolution). The anesthetized rats were secured during imaging in a custom-made tooth holder to stabilize their head position. The fluorescence signal was determined in a region of interest (ROI) placed at skull landmark lambda as a function of time, and quantified relative to the initial fluorescence intensity obtained at 0.5 s in a blinded manner. A white field image of the rat head was captured at the end of imaging prior to visualizing the lateral ventricles of the isolated brain hemispheres to verify bilateral carboxylate staining. Data analyses were performed in Image Studio 5.2 (LI-COR Biosciences – GmbH, Nebraska, US).

### Radioisotope flux assays

Isolated rat brains were kept in cold HEPES-aCSF (4°C, pH 7.35) for 10min prior to isolation of the lateral choroid plexuses. The isolated lateral choroid plexuses were subsequently placed in HEPES-aCSF (pH 7.56, 37°C) for 10min. The experiments were initiated by choroidal isotope accumulation (1 μCi ml^-1 86^Rb^+^, 022-105721-00321-0001, POLATOM, as a marker for K^+^ transport, and 4 μCi ml^-1 3^H-mannitol as an extracellular marker, PerkinElmer) for 2min (influx) or 10min (efflux). The influx experiments were conducted either in the presence of 2 mM ouabain (O3125, Sigma), 200 µM AZE, or in the appropriate vehicle, after which the choroid plexus was swiftly rinsed in cold isotope-free HEPES-aCSF (4°C ) and transferred to a scintillation vial. For the efflux experimentation, the choroid plexus was briefly washed (15 s) in 37°C HEPES-aCSF following the isotope accumulation step, and subsequentially transferred (at 20 s intervals) to different HEPES-aCSF solutions (37°C) containing either 200 μM AZE, 20 µM bumetanide (B3023, Sigma), or appropriate vehicle. The efflux medium from each of the solutions was transferred into separate scintillation vials. For both influx and efflux assays, the choroid plexus was dissolved in 100 µl Solvable (6NE9100, Perkin Elmer) prior to addition of 500 μl of Ultima Gold^TM^ XR scintillation liquid (6012119, PerkinElmer) and subsequent quantification in a Tri-Carb 2900TR Liquid Scintillation Analyzer (Packard). The ^86^Rb^+^ counts were corrected for ^3^H mannitol counts (extracellular background), and the natural logarithm of the choroid plexus content A_t_/A_0_ was plotted against time (58) to obtain the ^86^Rb^+^ efflux rate (s^-1^) by linear regression analysis.

### Intracellular Na^+^ measurement

Isolated lateral choroid plexuses were transferred into a recording chamber and perfused with HCO_3_ aCSF at room temperature (22 ± 1°C) for about 20min. Subsequently, the Na^+^-sensitive fluorescent dye SBFI-AM (sodium-binding benzofuran isophthalate acetoxymethyol ester, 2021E, ION Biosciences, dissolved in 20% Pluronic, F127) was pressure-injected into several regions of the choroid plexus, using a fine-tipped glass micropipette coupled to a pressure application system (PDES nxh, npi electronic, Tamm, Germany). The tissue was subsequently perfused with HCO_3_ -aCSF (~45min) to wash out excess dye and allow for de-esterification. Wide-field Na^+^ imaging was obtained with a variable scan digital imaging system (Nikon NIS-Elements v4.3, Nikon GmbH) coupled to an upright microscope (Nikon Eclipsle FN-PT, Nikon GmbH). The microscope was equipped with a ×40/N.A. 0.8 LUMPlanFI water immersion objective (Olympus Deutschland GmbH) and an orca FLASH V2 camera (Hamamatsu Photonics Deutschland GmbH). SBFI was alternately excited at 340 and 380 nm, and emission collected > 440 nm with a sampling rate of 0.5 Hz. Fluorescence emission was recorded from defined regions of interest (ROI) representing single cells, for 2min in HCO_3_ aCSF under baseline conditions, followed by perfusion with HCO_3_ -aCSF containing 200 µM AZE or 0.02% DMSO (control) and imaging for another 30min. SBFI signals were analyzed with OriginPro Software (OriginLab Corporation v.9.0). Background-correction was carried out for each ROI to obtain the fluorescence ratio (F340/F380). Linear regression analyses were performed on control and drug periods in a blinded fashion.

### RNAseq

Choroid plexus (lateral and 4^th^) were isolated from 5 AZE-treated and 5 control rats, pooled respectively, and stored in RNAlater® at -80°C. The RNA extraction and library preparation were performed by Novogene Company Limited, UK with NEB Next® Ultra™ RNA Library Prep Kit (NEB, USA) prior to their RNA sequencing (paired-end 150 bp, with 12 Gb output) on an Illumina NovaSeq 6000 (Illumina, USA). Program parameter settings for library build, mapping, and quantification, together with scripts for the gene annotation and analysis can be found at https://github.com/Sorennorge/MacAulayLab-RNAseq2-Acetazolamide. The 150 base paired-end reads were mapped to Reference genome Rnor_6.0 (Rattus_norvegicus v.103), only including protein coding genes (biotype), using Spliced Transcripts Alignment to a Reference (STAR) RNA-seq aligner (v 2.7.2a) (59). The mapped alignment generated by STAR were normalized to transcripts per million (TPM) (60) with RSEM (v. 1.3.3). The RNA sequencing data for human choroid plexus was obtained from Rodríguez-Lorenzo, (Geo: GSE137619, SRR10134643-SRR10134648) (61), and the RNA sequencing data for mouse choroid plexus was obtained from Lun et al. (Geo: GSE66312, SRR1819706-SRR18197014) (62). All human and mouse samples were quality checked with fastqc (63), and then trimmed with Trimmomatic (64) (Slidingwindow:4:20, and minimum length of 35bp). The human and mouse samples were mapped to reference human genome (Homo sapiens GRCh38.104) and mouse reference genome (Mus musculus GRCm39.104), both only including protein coding genes (biotype), with STAR (v 2.7.2a). The mapped alignment were quantified to TPM using RSEM (v. 1.3.3) and the mean from the human and the mouse samples were used for further analysis.

### Ventricular-cisternal perfusion

Rats were anesthetized, ventilated, and an infusion cannula (Brain infusion kit 2, Alzet) was stereotaxically placed in the right lateral ventricle (as described in ICP measurements), through which a pre-heated (37°C, SF-28, Warner Instruments) HCO_3_ -aCSF containing 1 mg ml^-1^ TRITC-dextran (tetramethylrhodamine isothiocyanate-dextran, MW = 150,000; T1287, Sigma) was infused at 9 μl min^−1^. CSF was sampled from cisterna magna at 5min intervals with a glass capillary (30-0067, Harvard Apparatus pulled by a Brown Micropipette puller, Model P-97, Sutter Instruments) placed at a 5° angle (7.5 mm distal to the occipital bone and 1.5 mm lateral to the muscle-midline). The fluorescent content of CSF outflow was measured in triplicate on a microplate photometer (545 nm, SyneryTM Neo2 Multi-mode Microplate Reader; BioTek Instruments), and the CSF secretion rate was calculated from the equation:

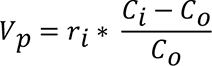

Where *V_p_* = CSF secretion rate (µl min^-1^), *r_i_* = infusion rate (µl min^-1^), *C_i_* = fluorescence of inflow solution, *Co* = fluorescence of outflow solution. The ventricles were perfused for 80min, and the production rate over the last 20min was used to calculate the average CSF secretion rate for the animal.

### Brain water content

Brains were swiftly isolated from euthanized rats, placed in a pre-weighed porcelain evaporation beaker (olfactory bulbs and medulla oblongata discarded), and immediately weighed. The brain was homogenized in the beaker with a steel pestle prior to oven-drying at 100°C for 3 days and subsequent determination of dry brain weights. Brain water contents were calculated from evaporated water, and expressed in ml water per g dry brain weight.

### ICP and MAP monitoring in awake rats

KAHA Sciences rat dual pressure telemetric system was implemented for monitoring ICP and MAP in non-anesthetized, awake rats. The implantation was performed as described in (65). Briefly, animals were given 5 mg kg^-1^ Caprofen (Norodyl Vet, Norbrook), 0.05 mg kg^-1^ buprenorphine (Tamgesic, Indivior), and 200 + 40 mg kg^-1^ sulfadiazin and trimethoprim (Borgal Vet, Ceva) s.c. prior to the surgery and 2 days post-surgery. The incision areas were shaved and sterilized with 0.5% chlorhexidine (Medic). An abdominal midline incision was made, the abdominal aorta isolated, and the first pressure probe was inserted into the aorta using a bent 23G needle. It was secured with tissue adhesive (Histoacryl, Enbucrilate; B. Braun) and surgical mesh (PETKM2002, SurgicalMesh; Textile Development Associates). The body of the telemetric device was secured to the abdominal wall, and 4-0 absorbable Vicryl suture (Ethicon) was used to close the abdominal muscles. The protruding second pressure probe was tunneled to the base of the skull using a 6 mm diameter stainless steel straw (Ecostrawz). The animal was placed into the stereotactic frame (Harvard apparatus), and the skull was exposed. Using 1.2 mm burr bits, two holes were drilled on the contralateral sides of the skull posterior to bregma. Stainless steel screws (00-96×3/32, Bilaney Consultants GmbH) were inserted into these holes, and served as anchors for stabilizing the system. The second pressure probe was placed epidurally in a third 1.4 mm drill hole placed between the two screws. The hole was subsequently filled with spongostan (Ethicon), and the probe was secured using surgical mesh and tissue adhesive. Dental impression material (Take 1 Advanced, Kerr) was applied over the catheter and the screw, and the skin incision was closed with non-absorbable 4-0 EthilonII suture (Ethicon), which was removed 10 days post-surgery. Animals were placed in their cages on the TR181 Smart Pads (Kaha Sciences), and data acquisition obtained at 1 kHz with PowerLab and LabChart software (v8.0, ADInstruments). Data was extracted from LabChart as 6min average values and outlying data points identified with GraphPad Prism (GraphPad Software). During the experimental series with 1× day AZE treatment, day 5 was discarded from the analysis due to the weekly cage change, which caused a visible disturbance in all measured parameters. The 3× day dosing experiments were performed on same animals that received 1× day dosing seven days after recovery from the previous treatment.

### Statistics

All data are presented as average ± SEM. Statistical significance analysis was performed with GraphPad Prism (GraphPad Software), and P ≤ 0.05 was considered statistically significant. 1way ANOVA with Tukey’s multiple comparison post hoc test was used in ICP and MAP analyses of nephrectomised, hyperventilated and i.c.v.-infused animals to include comparison with naïve animals. 2way ANOVA analysis was used in the blood gas analysis, and additional post-hoc analysis with Bonferroni’s (intact) or Tukey’s (nephrectomised, hyperventilated and i.c.v. infused) multiple comparisons test was performed for pCO_2_ data. 2way ANOVA with Bonferroni’s multiple comparisons test was also used to analyze the daily 2h average patterns in the telemetry experiments. Paired t-test was used to compare the telemetrically measured raw ICP values before and after 3x daily treatment. Unpaired t-tests were used in the rest of the analyses. Significance is represented as asterisks above the bar graphs and represent P > 0.05 = ns, P ≤ 0.05 = *, P ≤ 0.01 = **, P ≤ 0.001 = ***.

### Study approval

All animal experiments were performed in accordance to the European legislations for the care and use of laboratory animals, and approved by the Danish Animal Experiments Inspectorate (License no. 2016-15-0201-00944 and 2018-15-0201-01595) or the Animal Welfare Office at the Animal Care and Use Facility of the Heinrich Heine University Düsseldorf (institutional act no. O52/05).

## Results

### AZE effectively lowers ICP

To determine the modulatory effect of AZE on ICP in anesthetized and ventilated rats, AZE was delivered systemically as a bolus i.v. injection during simultaneous recording of the ICP. The base ICP of the experimental rats was 4.2 ± 0.1 mmHg, n = 34. We observed an abrupt spike in ICP immediately after AZE administration (to 133 ± 6%, n = 5) with a subsequent gradual decrease of the ICP to -48 ± 4% below the baseline level 2 h after the injection, n = 5 (Figure 1A-B). The control animals (bolus injection of vehicle) did not experience the initial ICP peak (103 ± 1%, n = 5), and the subsequent time-dependent decline in ICP amounted to -25 ± 4% below the baseline level, n = 5 (Figure 1A-B), which was significantly less than that of the AZE-treated rats (P < 0.01). AZE is recognized for its lowering effect on blood pressure (49), which in turn could affect ICP and/or CSF secretion (50). We therefore, in parallel with the ICP measurements, performed recordings of the mean arterial pressure (MAP). The initial MAP was 71.7 ± 1.3 mmHg, n = 34, amongst all the tested rats. The MAP remained stable for the duration of the experiment in both experimental animal groups (−5 ± 2% for AZE treated rats, and -6 ± 3% below baseline for control rats, n = 5, P = 0.9, Figure 1C-D). AZE thus exerts its effect on the ICP in a manner independent of the MAP.

**Figure 1.**
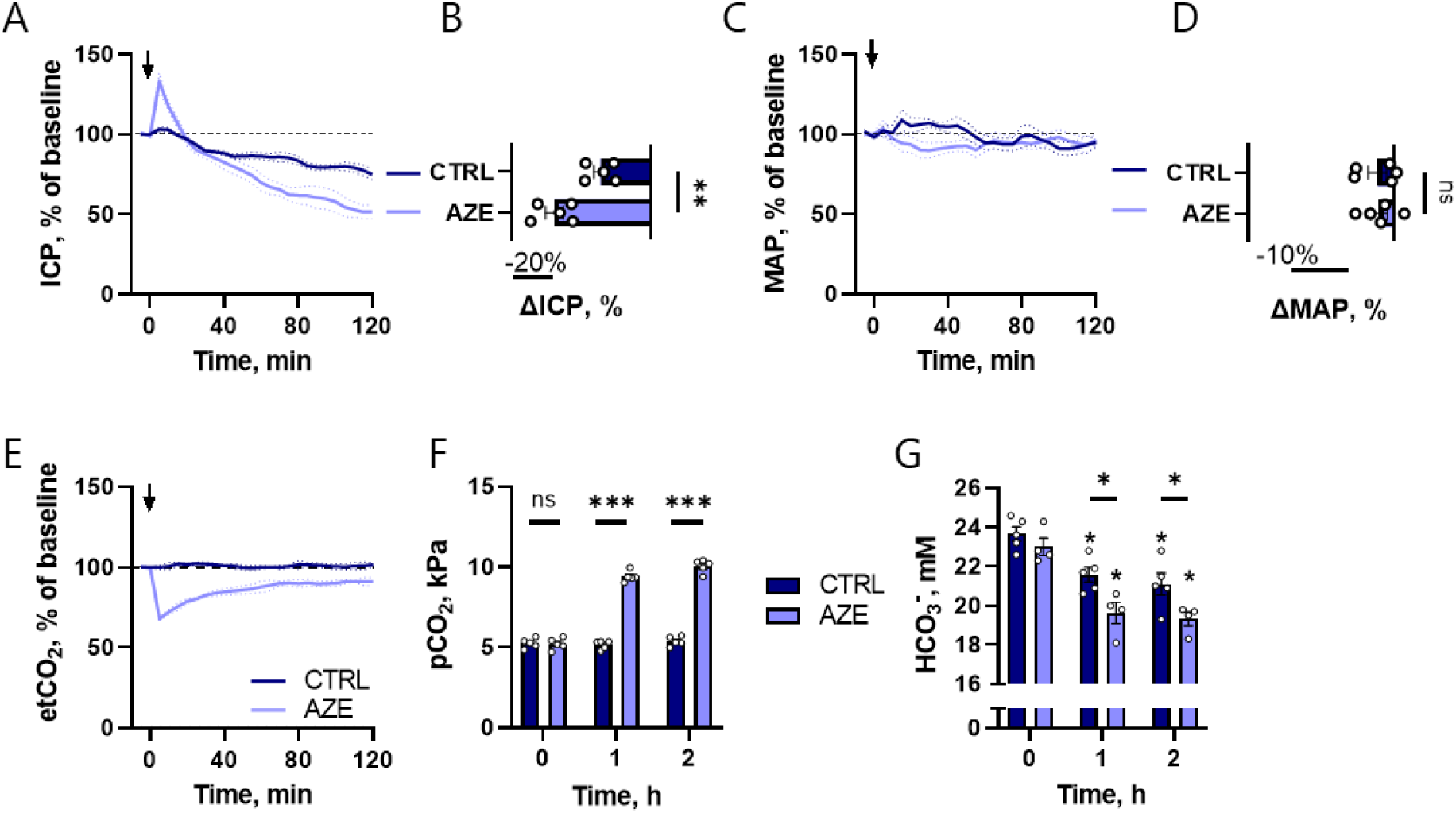
Effect of i.v. administration of 100 mg kg^-1^ AZE in anesthetized and ventilated rats. The ICP (**A**), MAP (**C**) and etCO_2_ (**E**) is presented as 5min average values normalized to the baseline (10min average value obtained before control solution or AZE administration) as a function of time. The change from baseline at 2h after the injection is shown for ICP (**B**) and MAP (**D**). Blood gas analysis was performed before (0h) and 1 and 2h after the injection, and results are presented for pCO_2_ (**F**) and for blood HCO_3_ , asterisks indicate significance for time variable, whereas bars indicate significance for treatment (**G**). Arrow indicates time of i.v. injection.

### AZE affects the systemic acid-base balance

As an inhibitor of the carbonic anhydrases, AZE affects the CO_2_–HCO_3_ conversion in the erythrocytes, and systemic delivery of AZE may therefore affect the blood gases in a manner that could indirectly affect CSF secretion or other physiological processes. We therefore monitored the exhaled etCO_2_ of the anesthetized and ventilated experimental rats at a constant rate. In contrast to the control animals (100 ± 1%, n = 5), administration of AZE caused an abrupt decrease in etCO_2_ (by -32 ± 1%, n = 5, P < 0.001), which remained reduced (to -9 ± 1% below the baseline etCO_2_) at the termination of the experiment (Figure 1E). This shift in etCO_2_ was reflected in the blood pCO_2_, which remained stable in control animals but increased dramatically in rats treated with AZE (from 5.2 ± 0.2 kPa to 10.1 ± 0.2 kPa 2h after AZE treatment, n = 5, P < 0.001, Figure 1F). Consequently, the blood pH decreased in AZE animals compared to controls (see all blood gas analyses in Supplementary Table S1). Taken together, the decreased exhaled etCO_2_, increased blood pCO_2_, and decreased pH indicate a severe respiratory acidosis due to carbonic anhydrase inhibition in the pulmonary endothelium and the circulating erythrocytes. The blood HCO_3_ levels decreased in the AZE treated experimental rat group (from 23.0 ± 0.4 mM to 19.3 ± 0.3 mM, n = 4) to a larger extent (P < 0.05) than what was observed in the control group (from 23.7 ± 0.4 mM to 21.1 ± 0.6 mM, n = 5, Figure 1G), suggesting an accompanying metabolic acidosis. Systemic administration of AZE thus causes severe acid-base disturbances, which could indirectly affect ICP and/or CSF secretion.

### AZE exert its effect on ICP independently of the systemic water homeostasis

An AZE-induced change in systemic HCO_3_ levels is likely to occur via the inhibitory action of AZE on carbonic anhydrases in the kidney proximal tubule epithelium (47). Such a process will impair HCO_3_ reabsorption and induce diuresis (66), which could lead to ICP reduction. To resolve whether AZE indirectly exerts its effect on ICP via its action on the kidney, we determined the effect of AZE administration in nephrectomized rats (Figure 2A). AZE treatment reduced the ICP of nephrectomized rats by (−47 ± 3%, n = 4, Figure 2B), which is similar to what was observed in non-nephrectomized rats (compare with -48 ± 4%, Figure 1A). The control treatment caused similar ICP reductions in the rats whether or not the animals had undergone nephrectomy (compare -27 ± 6%, n = 4, Figure 2B with - 25 ± 4%, n = 5 in Figure 1A). Nephrectomy caused a similar decrease in MAP in all experimental rats, irrespective of AZE administration (−30 ± 9% in the control group, and -31 ± 5% in the AZE group, n = 4 of each, P = 0.9, Figure 2C-D). Likewise, AZE-induced changes in etCO_2_ and pCO_2_ were indistinguishable between nephrectomized and intact animals (Figure 2E-F, all blood gas analyses available in Supplementary Table S2). The baseline HCO_3_ levels were reduced in both groups of experimental animals (compare 23.4 ± 0.3 mM in intact animals, n = 9 to 21.6 ± 0.5 mM, n = 8 in nephrectomized animals, p < 0.01), probably due to absent HCO_3_ reabsorption by the nephrectomized kidney. The blood HCO_3_^-^ levels declined during the course of the experiments (as observed in the intact animals, Figure 1G), but with no AZE-induced changes in HCO_3_ reabsorption in the nephrectomized rats (Fig. 2G). Taken together, the results indicate that the effect of AZE on ICP occurs independently of its inhibitory action on kidney function (including its HCO_3_ handling) and thus the systemic water homeostasis.

**Figure 2.**
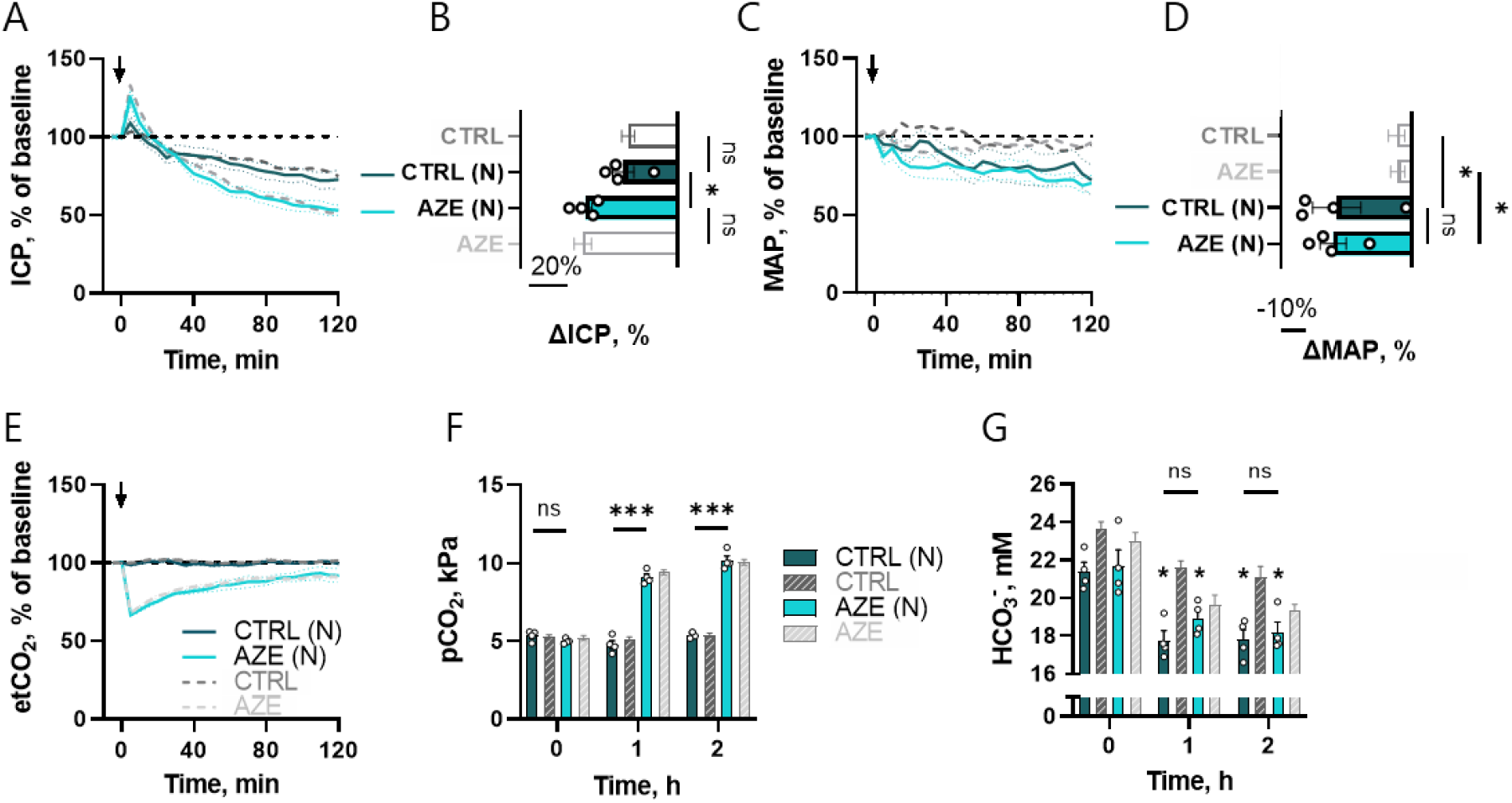
Effect of i.v. administration of 100 mg kg^-1^ AZE in anesthetized, ventilated and nephrectomized rats. ICP (**A**), MAP (**C**) and etCO_2_ (**E**) is presented in the same manner as Figure 1, as well as change at the 2h after the injection for ICP (**B**) and MAP (**D**). Results from the blood gas analysis are presented for pCO_2_ (**F**) and blood HCO_3_ , asterisks indicate significance for time variable, whereas bars indicate significance for treatment (**G**). Dark grey and light grey results are obtained from Figure 1. Arrow indicates time of i.v. injection. (N) = nephrectomized rats.

### Hyperventilation does not prevent AZE’s ability to lower ICP

To determine whether AZE exerts its effect on ICP in an indirect manner via altered CO_2_ dynamics in the blood, we attempted to prevent the AZE-induced CO_2_ retention in the blood by mechanical hyperventilation of the animals during the experimental procedure. Blood gas analysis (Supplementary Table S3) revealed that hyperventilation caused a 3.5 kPa reduction in pCO_2_ in control animals by the end of experiment (from 5.4 ± 0.1 kPa, n = 5 in intact animals to 1.9 ± 0.1 kPa, n = 4 in hyperventilated, P < 0.001), but a 1.6 kPa reduction in AZE-treated animals (from 10.1 ± 0.2 kPa, n = 5 in intact to 8.4 ± 0.3 kPa, n = 4 in hyperventilated, P < 0.05, Figure 3A). Hyperventilation thus did not revert the AZE-induced change in pCO_2_ to the level of the control animals (Figure 3A). Despite the partial prevention of the AZE-induced elevation of blood CO_2_ levels by hyperventilation, AZE treatment caused a reduction of ICP (−65 ± 3%, n = 4, Figure 3B-C) that was augmented compared to that obtained with conventional ventilation (−48 ± 4%, n = 5, Figure 1A, P < 0.05), yet the same was not observed in the hyperventilated control animals (compare -26 ± 4%, n = 4 Figure 3C with -25 ± 4%, n = 5 in Figure 1A). Hyperventilation reduced the MAP to a similar extent (P = 0.3) in both the control (−30 ± 1%, n = 4) and AZE (−24 ± 5%, n = 4) group (Figure 3E-F), supporting the finding from Figure 1D that AZE treatment did not reduce the MAP. Taken together, AZE-induced increase in blood pCO_2_ does not contribute to ICP reduction.

**Figure 3.**
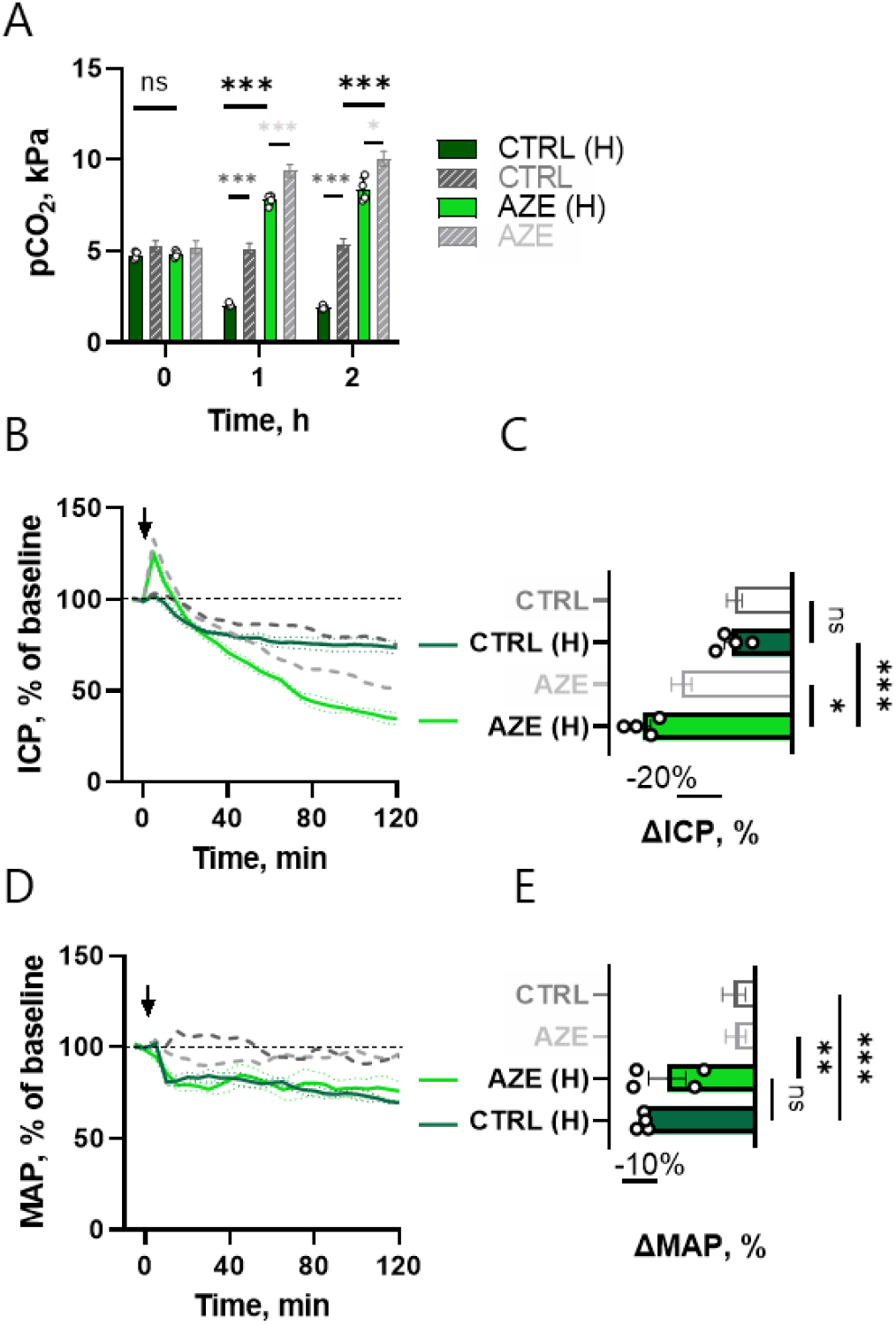
Effect of i.v. injection of 100 mg kg^-1^ AZE in anesthetized and hyperventilated rats. Hyperventilation was induced by increasing the tidal volume and the respiratory rate by 50% from the baseline value over 5min after the injection of treatment, which caused pCO_2_ reduction (**A**). The ICP (**B**) and MAP (**D**) are presented as function of time in the same manner as in Figure 1, with end 2h change shown in **C** (ICP) and **E** (MAP). Dark grey and light grey results are obtained from Figure 1. Arrow indicates time of i.v. injection. (H) = hyperventilated rats.

### Intraventricular AZE prevents systemic disturbances, but retains its ICP lowering effect

AZE thus facilitates a reduction in ICP independently of its effect on blood pressure, kidney function, and blood gas content. To obtain a scenario in which AZE could be administered without systemic disturbances, we applied AZE intracereboventricularly (i.c.v.). AZE application directly into the lateral ventricle of anesthetized rats did not disturb the MAP (Figure 4A-B), the blood pCO_2_ (Figure 4C), the blood HCO_3_ concentration (Figure 4D), or any other blood gas parameter including pH (Supplementary Table S4). Nevertheless, AZE delivery in this manner caused an ICP reduction (−51 ± 5%, n = 4), significantly larger than that of the control group (−16 ± 7%, n = 4, P < 0.01, Figure 4E-F), but similar to that obtained upon i.v. injection of the inhibitor (48 ± 4%, n = 5, see Figure 1A). In addition, the initial AZE-mediated ICP peak observed with systemic delivery of AZE (Figure 1A) was absent with the i.c.v. delivery of the inhibitor (Figure 4E). AZE administration directly into the brain thus lowers the ICP equally effectively as with systemic delivery, but independently of the AZE-mediated modulation of systemic parameters that could indirectly affect ICP. AZE therefore appears to serve its modulatory action on the ICP via a pathway residing in the brain tissue.

**Figure 4.**
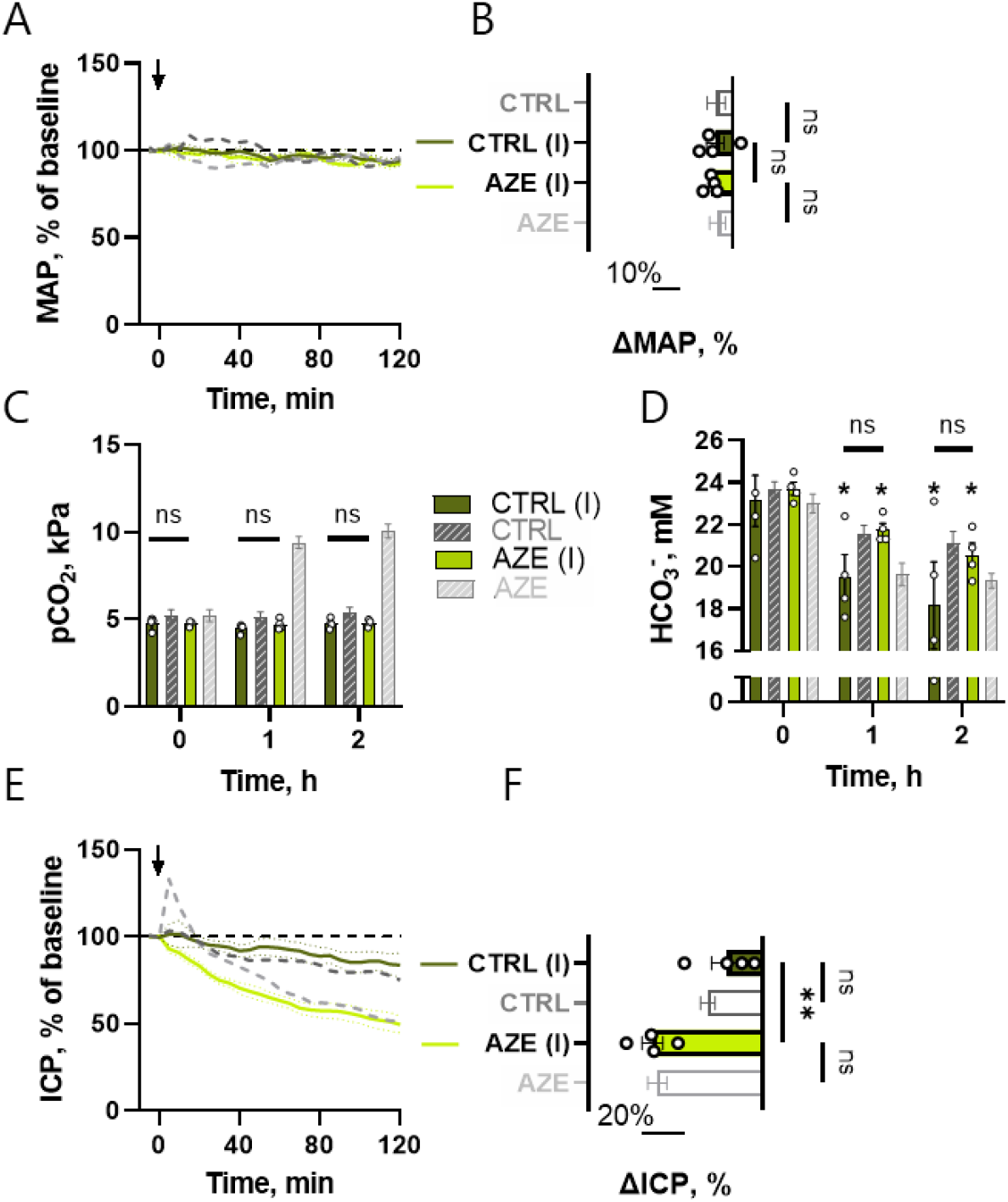
Effect of i.c.v. infusion of either control solution or 18 mM (expected ventricular concentration of max 500 µM) AZE at a rate of 0.5 µl min^-1^ in anesthetized and ventilated rats. MAP (**A**) is presented in the same manner as Figure 1, with 2h end MAP shown at **B** (ΔMAP_CTRL(I)_ = -6 ± 3%, ΔMAP_AZE(I)_ = -7 ± 1%, P = 0.9), as well as blood pCO_2_ (**C**) and blood HCO_3_ , asterisks indicate significance for time variable, whereas bars indicate significance for treatment (**D**). Effect of AZE i.c.v. as a function of time is shown in **E**, with end 2h change at **F**. Dark grey and light grey results are obtained from Figure 1. Arrow indicates the start of i.c.v. infusion. (I) = rats receiving i.c.v. delivery of AZE.

### AZE treatment lowers CSF flow in anesthetized rats

To determine if the AZE-mediated lowering of the ICP occurred by a reduction in the rate of CSF secretion, we assessed this parameter with a swift, minimally invasive approach based on imaging fluorescent dye flow in the ventricles of anesthetized rats (67) (Figure 5A). The fluorescent dye movement was reduced by 40% upon AZE treatment, whether the inhibitor was delivered i.c.v. (compare 0.16 ± 0.01 a.u. min^-1^, n = 4 with 0.10 ± 0.02 a.u. min^-1^, n = 4, P < 0.05, Figure 5B) or i.v. (compare 0.16 ± 0.02 a.u. min^-1^, n = 6 with 0.09 ± 0.01 a.u. min^-1^, n = 4, P < 0.05, Figure 5C). AZE thus exerts its effect on the ICP, at least in part, by reducing the CSF secretion rate.

**Figure 5.**
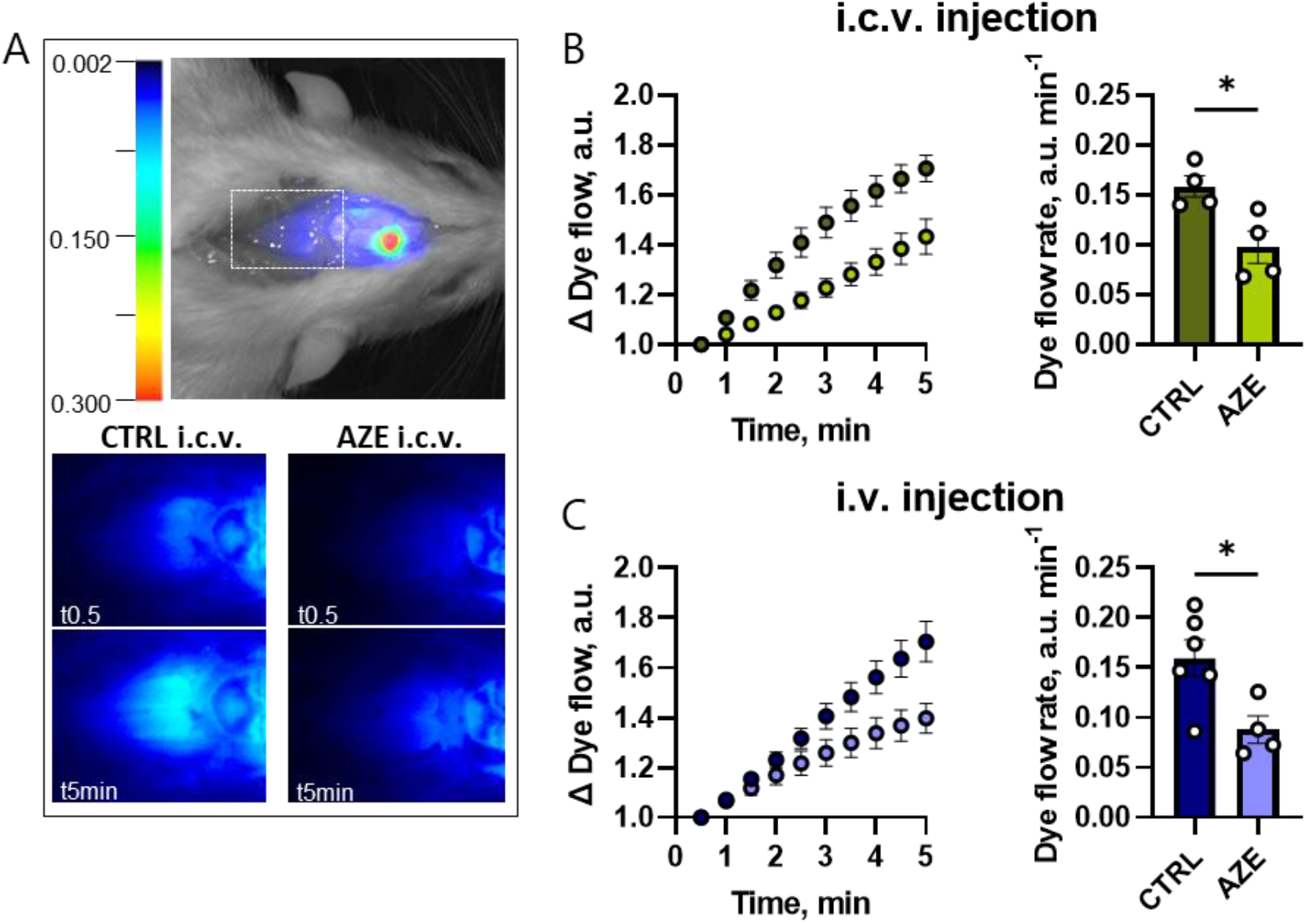
Live imaging of fluorescent dye as a proxy of CSF secretion following treatment with AZE. The top panel in **A** illustrates pseudo-color fluorescence superimposed on a white light image from a rat after ventricular injection of IRDye 800 CW carboxylate dye. The white box below lambda illustrates the area for dye intensity quantification. The bottom panel illustrates representative images of fluorescence signal 0.5 and 5min after i.c.v. injection of control solution or 2 mM AZE solutions (expected ventricular concentration 200 µM). Quantification of the dye flow normalized to the first image after i.c.v. injection is shown in panel **B**. Panel **C** shows the quantification of the dye flow normalized to the first image after i.v. injection of control solution or 100 mg kg^-1^ AZE.

### AZE does not affect the transport rate of Na^+^/K^+^-ATPase or NKCC1

AZE-dependent inhibition of carbonic anhydrases indirectly inhibits HCO_3_ transporters by reducing the available substrate, and most likely exerts its effect on CSF secretion by affecting some of these transport mechanisms located on both the sides of the CSF secreting choroid plexus epithelium (35). However, AZE could serve indirect effects on other choroidal transport mechanisms, such as its proposed action on the Na^+^/K^+^-ATPase (14). To determine a putative AZE-mediated reduction in transport rate of other key CSF secreting transporters, we performed *ex vivo* radioactive ^86^Rb^+^ efflux and influx assays, which serve as a functional read-out for the activity of NKCC1 and Na^+^/K^+^-ATPase, respectively. The NKCC1 activity was determined as the efflux rate of ^86^Rb^+^ (serving as a congener for the transported K^+^) that is sensitive to the NKCC1 inhibitor bumetanide (Figure 6A). The ^86^Rb^+^ efflux rate constant was diminished by ∼50% by inclusion of bumetanide (compare 0.33 ± 0.02 with 0.17 ± 0.01 min^-1^, n = 6 of each, P < 0.001, Figure 6A-B). Exposure to AZE did not affect the ^86^Rb^+^ efflux rate (compare 0.32 ± 0.02 in control with 0.34 ± 0.02 min^-1^ in AZE, n = 6 of each, P = 0.8, Figure 6C-D). Na^+^/K^+^-ATPase activity was monitored as the ouabain-sensitive influx of ^86^Rb^+^. Application of the selective Na^+^/K^+^-ATPase inhibitor oubain (2 mM) reduced the influx rate by 70% (compare 2772 ± 501 cpm in control with 794 ± 217 cpm in ouabain, n = 6 of each, P < 0.01, Figure 6E), whereas AZE did not alter the uptake rate (compare 2141 ± 119 in controlwith 2168 ± 361 cpm in AZE, n = 6 of each, P = 0.9, Figure 6F). These data indicate that the AZE-mediated reduction in the CSF secretion rate does not occur via indirect modulation of the NKCC1 or the Na^+^/K^+^-ATPase, and therefore must be via the various HCO_3_ transporters located in the choroid plexus epithelium.

**Figure 6.**
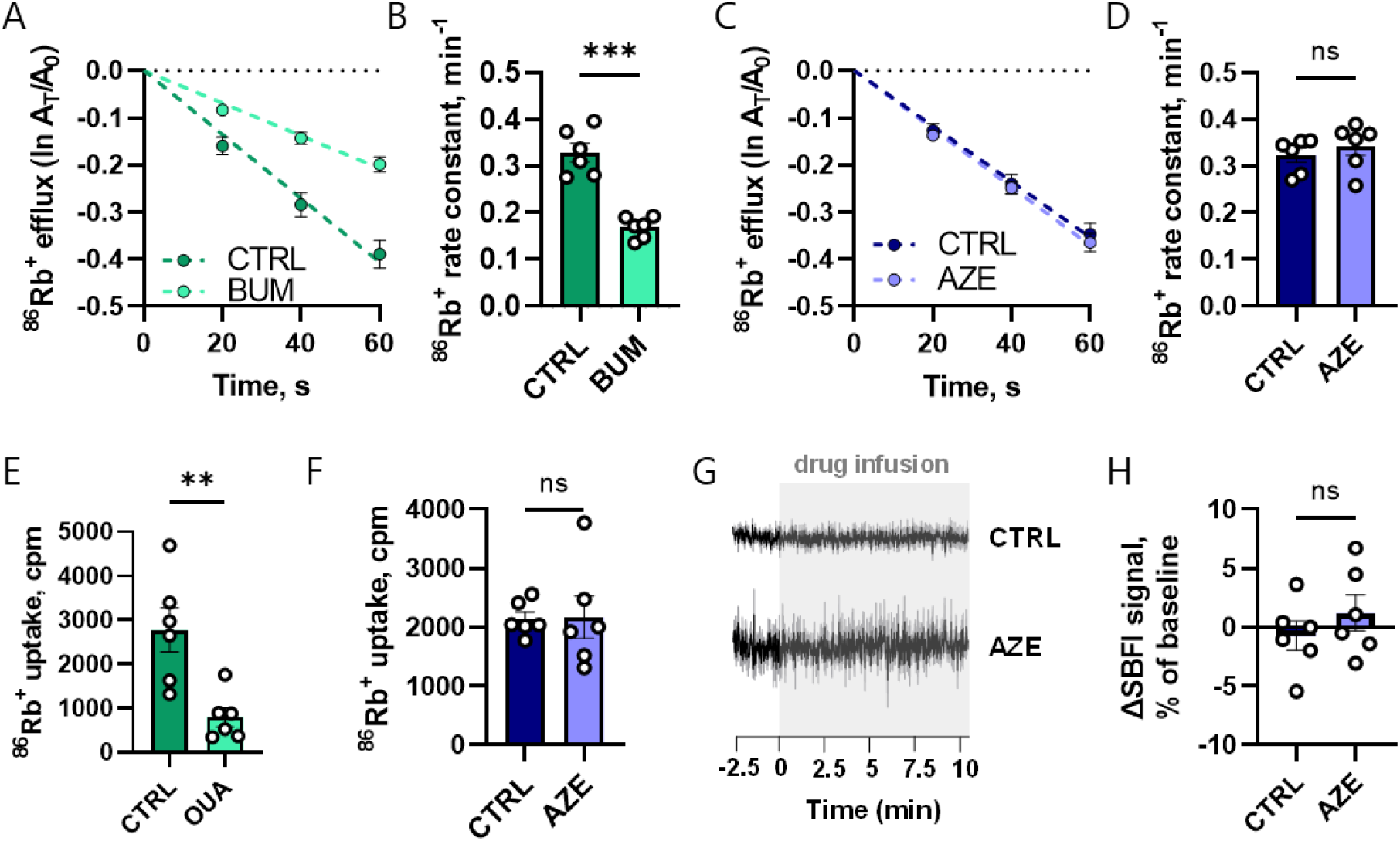
AZE’s effect on the activity of selected key CSF secreting transporters. ^86^Rb^+^ efflux rate (a read out for NKCC1 activity) is presented as a function of time upon treatment with either 20 µM bumetanide (**A**) or 200 µM AZE (**C**). The efflux rate constant was quantified using linear regression analysis for bumetanide (**B**) and AZE (**D**). ^86^Rb^+^ uptake rate (read out for Na^+^/K^+^-ATPase activity) is shown after treatment with 2 mM ouabain (**E**) and after treatment with 200 µM AZE (**F**). Na^+^i sensitive SBFI dye traces during exposure to 200 µM AZE or control solution are shown in **G**, and quantified in **H** (CTRL: -0.7 ± 1.2%, AZE: 1.2 ± 1.5%, n = 6, P = 0.34).

To reveal the choroidal [Na^+^]i dynamics upon AZE treatment, acutely isolated *ex vivo* choroid plexus tissue was loaded with the Na^+^-sensitive fluorescent dye SBFI, and monitored by wide-field SBFI imaging. The SBFI signal stayed stable during the AZE exposure (Figure 6G-H), suggesting that the [Na^+^]i remained undisturbed during AZE’s inhibitory action. These data support the notion that NKCC1 and the Na^+^/K^+^-ATPase activity occurred uninterrupted in the presence of AZE. Of note, an AZE-mediated reduction of the CSF secretion rate with no concomitant changes in [Na^+^]i could occur by inhibition of the various HCO_3_ transporters located on both epithelial membranes of the choroid plexus.

### AZE does not reduce choroidal expression of carbonic anhydrases or key transporters implicated in CSF secretion

To assess carbonic anhydrase expression in the rat choroid plexus, and to evaluate whether chronic AZE treatment caused changes in the functional properties of the choroid plexus related to CSF secretion, we performed RNAseq analysis on choroid plexus obtained from animals that received AZE (or control solution) once daily for 7 days. 13 different CA isoforms were expressed in the rat choroid plexus, with CA2 being expressed nearly one order of magnitude higher than the subsequent CA14 (Table 1). Comparison of the carbonic anhydrase isoform transcripts detected in rat choroid plexus with those obtained from mouse (62) and human (61) choroid plexus, revealed that seven were detected in all three species at moderate to high levels (CA2, CA14, CA12, CA11, CA13, CA5B, CA3 in order of abundance), which suggests their functional importance in the tissue. CA4 was detected in rat and human choroid plexus, whereas two isoforms (CA6, CA9) were found only in the rat choroid plexus. Three isoforms were below reliable detection level in the rat choroid plexus (CA1, CA7, CA8). Chronic AZE treatment did not cause a general down regulation of the choroidal CAs, although CA4 abundance was reduced to approximately half. In contrast, a compensatory *elevation* seemed to occur for transcripts encoding CA9 and CA12 , as well as for two of the key choroidal transport mechanisms, the Na^+^/K^+^-ATPase and NBCe2 (Table 2). Therefore, chronic exposure of AZE does not appear to provide a sustained reduction of the choroidal CSF secretion apparatus.

**Table 1.** Comparison of carbonic anhydrase expression levels in the control (administered p.o. saline once daily for 7 days) rat choroid plexus with the RNAseq data available from mice (62) and from humans (61). The isoforms are presented from the highest to the lowest expression in rats. TPM: transcript per million.

|  |  | **Rat,**<br>**TPM** | **Mouse,**<br>**TPM** | **Human,**<br>**TPM** |
| --- | --- | --- | --- | --- |
| 1 | CA2 | 785 | 368 | 1064 |
| 2 | CA14 | 92 | 243 | 50 |
| 3 | CA4 | 15 | - | 6 |
| 4 | CA12 | 13 | 828 | 107 |
| 5 | CA11 | 10 | 42 | 28 |
| 6 | CA6 | 8 | - | - |
| 7 | CA13 | 7 | 3 | 0.9 |
| 8 | CA9 | 3 | - | - |
| 9 | CA5B | 3 | 4 | 9 |
| 10 | CA3 | 0.6 | 4 | 1 |
| 11 | CA8 | 0.3 | 1 | 3 |
| 12 | CA7 | 0.2 | - | - |
| 13 | CA1 | 0.2 | - | 1 |
|  | CA10 | - | 132 | - |
|  | CA15 | - | 7 | - |

**Table 2.**
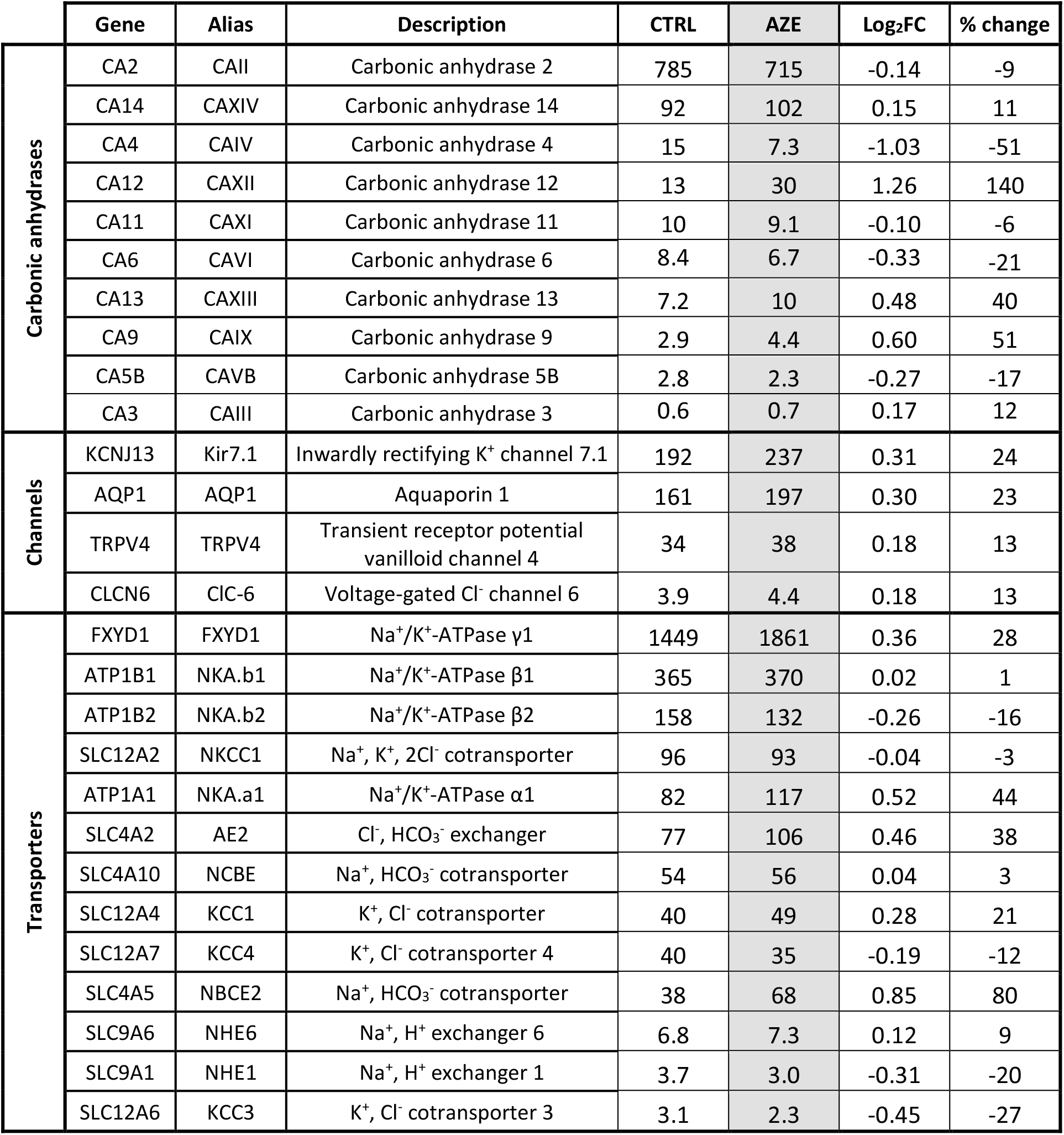
Expression levels, the fold change (Log_2_FC), and the % change of main transporters, channels and carbonic anhydrases in the rats choroid plexus after treatment with either control solution (CTRL) or 100 mg ml^-1^ AZE p.o. for 7 days (AZE), in transcripts per million (TPM). Control values for carbonic anhydrases are those from Table 1.

### Chronic exposure to AZE does not provide prolonged effects on the brain water content or CSF secretion rate

It is evident that acute intake of AZE reduces the ICP, which, at least partially, arises from AZE-mediated reduction in CSF secretion rate. However, the limited efficacy in the clinical setting called for an evaluation of AZE’s effect upon chronic intake. To determine whether prolonged exposure to AZE altered the CSF dynamics, experimental rats were treated with AZE (or control solution) once daily for 7 days. The brain water content obtained 24h after last treatment remained undisturbed following such treatment (compare 3.67 ± 0.03 ml g^-1^ dry brain weight in control animals to 3.69 ± 0.01 ml g^-1^ dry brain weight in animals treated with AZE, n = 5 of each, P = 0.5, Figure 7A). This notion of undisturbed fluid dynamics after chronic exposure to AZE was underscored by the similar CSF secretion rate obtained the day following completion of the treatment regime with the ventricular-cisternal perfusion assay (67) in treated animals (5.4 ± 0.3 μl min^-1^) versus control animals (4.7 ± 0.6 μl min^-1^, n = 4 of each, P = 0.4, Figure 7B-C). Chronic delivery of AZE (once daily) therefore does not appear to lead to a sustained (24h) reduction of brain fluid content or secretion thereof.

**Figure 7.**
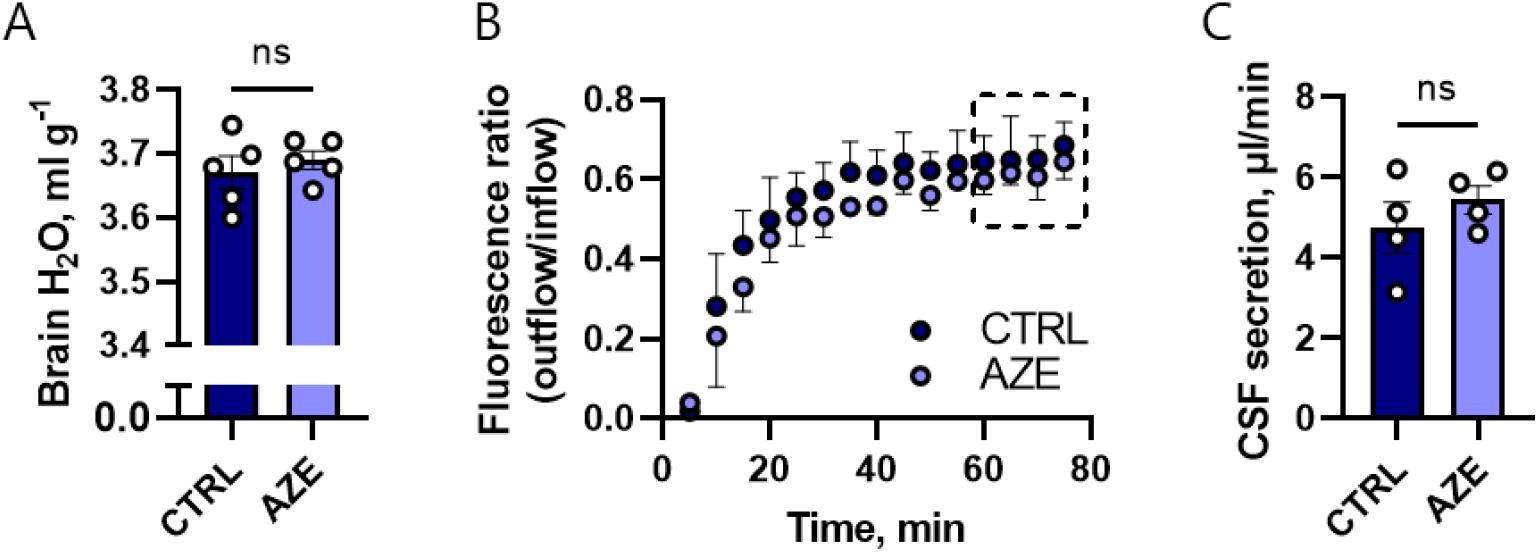
Effect of 1× daily administration of 100 mg kg^-1^ p.o. AZE or control solution on brain water content and CSF secretion rate, measured 24 h after the last dose. Panel **A** represents the brain water content expressed as ml g^-1^ dry weight. Figure **B** shows the average dextran dye dilution for control and AZE groups as a function of time. The dashed boxed indicates the period from which the CSF secretion rate was calculated, and the average value for this 20min period is presented in **C**.

### Chronically administered AZE has a short-lived effect on ICP

To monitor the physiological impact of AZE treatment as a function of time in awake and freely moving rats, we monitored the ICP and MAP simultaneously during AZE administration by employing Kaha telemetric dual pressure transmitters. Following implantation of the device, the ICP declined from 6.0 ± 0.7 mmHg on 1^st^ day post-surgery to a stable level around 3.5 ± 0.3 mmHg 10 days post-surgery, n = 8 (Figure 8A), a pattern that was mimicked by the heart rate (from 413 ± 7 bpm to 338 ± 9 bpm, n = 8, Figure 8B). The MAP was stable during this recovery time (91 ± 3 mmHg post-surgery, and 94 ± 2 mmHg on day 10, n = 8, Figure 8C), but day-night time oscillations were not visible until 7 days post-surgery, confirming that a recovery period of at least 7 days is required prior to collection of physiological data.

**Figure 8.**
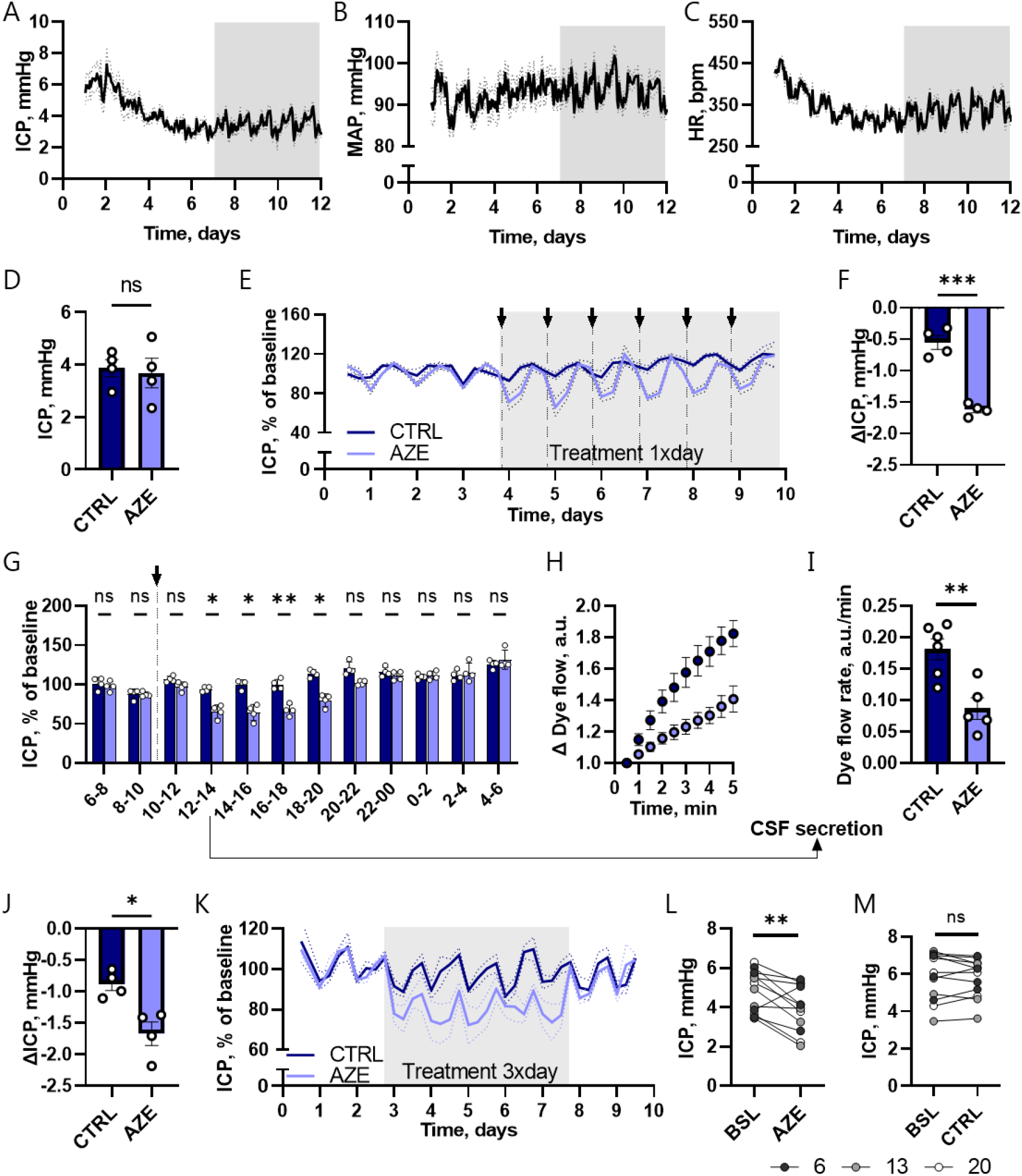
Telemetric measurements of ICP in freely moving awake rats before and during p.o treatment with 100 mg kg^-1^ AZE. Top panels represent the recovery period after implantation of telemetric device and show daily fluctuations in ICP (**A**), MAP (**B**) and heart rate (**C**). Panel **D** shows the average ICP over 3 days (72h) before initiation of treatment (ICP_CTRL_ = 3.9 ± 0.3 mmHg, ICP_AZE_ = 3.7 ± 0.6 mmHg, n = 4 in each, P = 0.8). The % change in ICP, normalized to the average 24h ICP (baseline) before treatment initiation, is shown in **E** for 1× daily p.o. delivery of 100 mg kg^-1^ AZE or control solution. The difference between the lowest 1h average ICP value (within the first 7h after AZE or control solution injection) and the baseline ICP is shown in **F**. The ICP fluctuations over the 24h period after drug administration are represented in **G**, where an average ICP for each 2h period was calculated for controls and AZE treated rats across the six treatment days. 2-3h after p.o. drug administration an IRDye 800 CW carboxylate dye was injected intraventricularly, and the dye flow (**H**) and rate (**I**) was used as a proxy to assess the CSF secretion rate. The maximum effect of the 3× day treatment with 100 mg kg^-1^ AZE or control solution for 5 days is shown in panel **J**. The ΔICP was calculated as difference between the baseline ICP and minimum 1 h average ICP during the 7h after each (of 14) AZE or control solution p.o. delivery. In **K**, ICP fluctuations during the 3× day treatment, normalized to baseline ICP are shown. The average ICP at 6, 13 and 20 o’clock (the hour before p.o. treatment administration) was calculated for the baseline period, and compared to average ICP at the same time points during 3× day treatment with 100 mg kg^-1^ AZE (**L**) or control solution (**M**).

At the termination of the recovery period, the 72h average ICP for the experimental rats was 3.8 ± 0.4 mmHg, n = 8, and not significantly different between the two experimental groups, P = 0.8 (Figure 8D). The treatment period was initiated by p.o. administration of AZE (or control solution) and repeated every 24h for 6 days, during which the ICP and MAP was monitored continually. The ICP pattern observed in the control rats resembled that of the rats prior to initiation of the treatment regime (Supplementary Figure 1A). The ICP fluctuated with the diurnal cycle, but deflected more in rats exposed to AZE compared to the control rats (Figure 8E) with a significantly higher AZE-induced ICP reduction (−1.6 ± 0.1 mmHg, n = 4) than that observed in the control group (−0.5 ± 0.1 mmHg, n = 4, P < 0.001), Figure 8F. The AZE-mediated ICP-reduction lasted for approximately 10h post-treatment (Figure 8G), after which the ICP of the AZE-treated rats matched that of the control rats (compare 111 ± 3% of baseline, n = 4 for AZE rats at 10-12h post-treatment with 116 ± 3% of baseline, n = 4, P = 0.2). An identical pattern was observed with the subsequent AZE administration the following days (Figure 8G).

To determine whether the AZE-mediated ICP deflection originated from a reduced rate of CSF secretion, as observed in the acutely treated animal experimental protocols, we measured the CSF secretion rate with the fluorescent imaging technique employed in Figure 5A. This swift protocol allowed us to resolve the CSF secretion rate at exactly 2-3h after the last of the six AZE (or control solution) doses was administered. The rate of CSF secretion was reduced 50% compared to that obtained in the control rats (compare 0.18 ± 0.02 a.u. min^-1^, n = 6 with 0.09 ± 0.02 a.u. min^-1^, n = 5, P < 0.01, Figure 8H-I). These data support the notion that AZE-mediated reduction in the CSF secretion rate underlies its modulatory effect on ICP.

The AZE-mediated reduction in ICP was not mirrored in change of MAP or heart rate. The MAP of the experimental rats was 93 ± 2 mmHg (n = 8, Supplementary Figure 1B) prior to initiation of the treatment period, and were indistinguishable between the AZE and the control group throughout the experimental period (Supplementary Figure 1C). Same was observed for the heart rate, with the baseline heart rate of 343 ± 12 bpm (n = 8, Supplementary Figure 1D), and no AZE-mediated effect on this parameter (Supplementary Figure 1E). These data suggest that AZE effectively reduces the ICP in awake and freely moving rats in a manner independent of cardiovascular effects.

To resolve whether a frequent dosing regimen can provide the sustained decrease in ICP required for clinical efficacy of such pharmacological treatment, the rats were dosed 3× daily at intervals of 7-7-10h. The maximal AZE-mediated ICP reduction was similar to that obtained with a single daily dose (compare -1.7 ± 0.2 mmHg for 3× day treatment (Figure 8J) with -1.6 ± 0.1 mmHg for 1× treated animals (Figure 8F), n = 4 in each group, P = 0.8). The 3× daily dosing, however, provided a sustained decrease in ICP (Figure 8K), which at no point reached the baseline levels (compare 4.8 ± 0.3 mmHg baseline ICP with 3.9 ± 0.3 mmHg ICP just prior to any of the next AZE treatments, n = 12 (4 biological replicates at 3 different time points) in each of the three groups, P < 0.01, Figure 8L). In contrast, the control group ICP remained at baseline throughout the experiment (compare 5.8 ± 0.4 mmHg baseline ICP with 5.8 ± 0.3 mmHg before each treatment with control solution, n = 4 in each group, P = 0.7). Regular administration with AZE during the day thus ensures a sustained ICP reduction of the experimental rats.

## Discussion

Here, we provide evidence for AZE’s ability to reduce ICP in an in vivo rodent model, and showcase that this outcome arises from AZE’s *direct* action on the CSF secretory machinery, and not via modulation of other physiological processes that may *indirectly* affect CSF secretion. We demonstrate that AZE’s ability to lower ICP and reduce the rate of CSF secretion is short lasting, and thus confirm the necessity of frequent dosing in the clinical setting.

AZE has been demonstrated to reduce the CSF secretion rate in various species (16–31), irrespective of its route of delivery (i.v. or i.c.v.), as AZE can cross the cell membrane and reach the choroidal carbonic anhydrases. The present study confirmed that AZE reduces the CSF secretion rate to a similar extent (40%) whether administered i.v., i.c.v., or p.o. The AZE-mediated reduction in CSF secretion did not occur via an effect on the transport activity of the Na^+^/K^+^-ATPase or the NKCC1, which are both involved in the CSF secretion (21, 22, 67), but AZE most likely acts via its effect on various HCO_3_ transporters localized in the choroid plexus and implicated in CSF secretion (35). However, it remained unresolved if such modulation of the CSF secretion rate is directly represented in a change in ICP and/or ventricular volume. The latter two parameters do not necessarily go hand in hand, as exemplified in idiopathic intracranial hypertension, in which the ICP is elevated without enlarged ventricles (12), and, in contrast, normal pressure hydrocephalus, in which the ventricles are enlarged, but an ICP elevation is absent or minor (68). The etiology of these diseases is not fully understood, but AZE treatment may still be employed to treat the symptoms in these patient groups (12, 69), despite this approach being questioned by several clinical trials (8, 9).

We demonstrate here that delivery of AZE to healthy experimental rats led to a 40% reduction of their ICP, irrespective of the route of administration (i.v., i.c.v., or p.o. by gavage), and of whether the animals were under anesthesia or freely moving. This finding is in line with earlier studies that validated AZE’s efficacy in lowering the ICP (13, 14), but contrasts a study on sedated rats in which no difference in ICP was observed upon AZE delivery either s.c. or p.o. (in Nutella), when controlling for solution osmolarity (15). The route of AZE administration and/or the lack of mechanical ventilation during the experimental procedure may have caused the difference in observations. In an animal model of hemorrhagic stroke, AZE was shown to prevent ICP spikes, without reducing the average ICP (70), whereas in a rat model of intracerebral haemorrhage, AZE reduced the brain water content and improved the functional outcome (71).

In the current study we demonstrated that a significant decrease in ICP is observed >2 h after p.o. AZE administration in awake rats – a comparable effect to that observed in patients (72). CSF secretion measurements at this exact time point revealed an underlying robust reduction in CSF secretion, which aligned well with that observed upon acute delivery of AZE (i.v. or i.c.v.) in anesthetized rats. Telemetric ICP measurements provided important insight into AZE’s mode of action, which could be employed for designing clinical treatment regimens. Reliable and stable ICP measurements in awake rats were observed seven days post-surgery. A daily single dose of AZE, equivalent to a clinically employed 1 g single dose in humans (12, 15), was effective in lowering the ICP. Yet, the ICP returned to baseline within 10-12 h after the treatment. This lack of a prolonged effect of repeated single daily AZE doses was reflected in identical brain water content in the two experimental groups and an undisturbed rate of CSF secretion obtained the day after the final dosing, and supported by lack of downregulation of choroidal transcripts encoding carbonic anhydrase or those encoding transport proteins involved in CSF secretion. Delivery of AZE at regular intervals 3× daily, rather than once daily, led to a significant decrease of the ICP throughout the 24h cycle, although upon discontinued treatment, the ICP returned to baseline levels. This finding suggests that frequent dosing and strict patient compliance are crucial for effective symptomatic relief of elevated ICP via treatment with AZE.

The AZE-mediated reduction in ICP appears to be a direct result of reduced CSF secretion. Such reduction of CSF secretion can occur directly by inhibition of the carbonic anhydrases in the choroid plexus. We detected expression of the CA isoforms 2, 14, 4, 12, 11, 6, 13, 9, 5B, 3 (in order of expression level) in the rat choroid plexus (Figure 9), the majority of which are detected at the transcript level in mouse and human choroid plexus as well (except CA6, CA9). Some of these choroidal CA isoforms (CA2, CA3, CA9, CA12, CA14) have been verified at the protein level (42–46), but their individual quantitative contribution to CSF secretion remains unresolved. AZE-mediated reduction of ICP could, however, occur indirectly by affecting carbonic anhydrases in other tissues and cell types in the body. In this manner, AZE has been proposed to affect blood pressure (48, 49), which could reduce blood flow to the choroid plexus (23, 73), and lead to a decrease in the CSF secretion rate (50). Despite the robust AZE-induced ICP reduction, we did not detect an AZE-mediated decline in the MAP whether delivered i.v. or i.c.v. to anaesthetized rats or p.o. to freely moving rats with telemetric monitoring of their MAP. AZE administration reduced blood [HCO_3_ ], which, in itself, could affect the transport rate of the choroidal HCO_3_ transporters supporting the CSF secretion (40), and thus indirectly lower the CSF secretion. This AZE-mediated [HCO_3_ ]_blood_ modulation was, however, abolished with i.c.v. delivery of AZE or upon functional nephrectomy of the experimental rats prior to i.v. delivery of AZE, while the ICP reduction endured in both of these experimental paradigms. The ability of AZE to reduce ICP via a decrease of the rate of CSF secretion thus occurred independently of its potential effect on blood pressure and kidney function.

**Figure 9.**
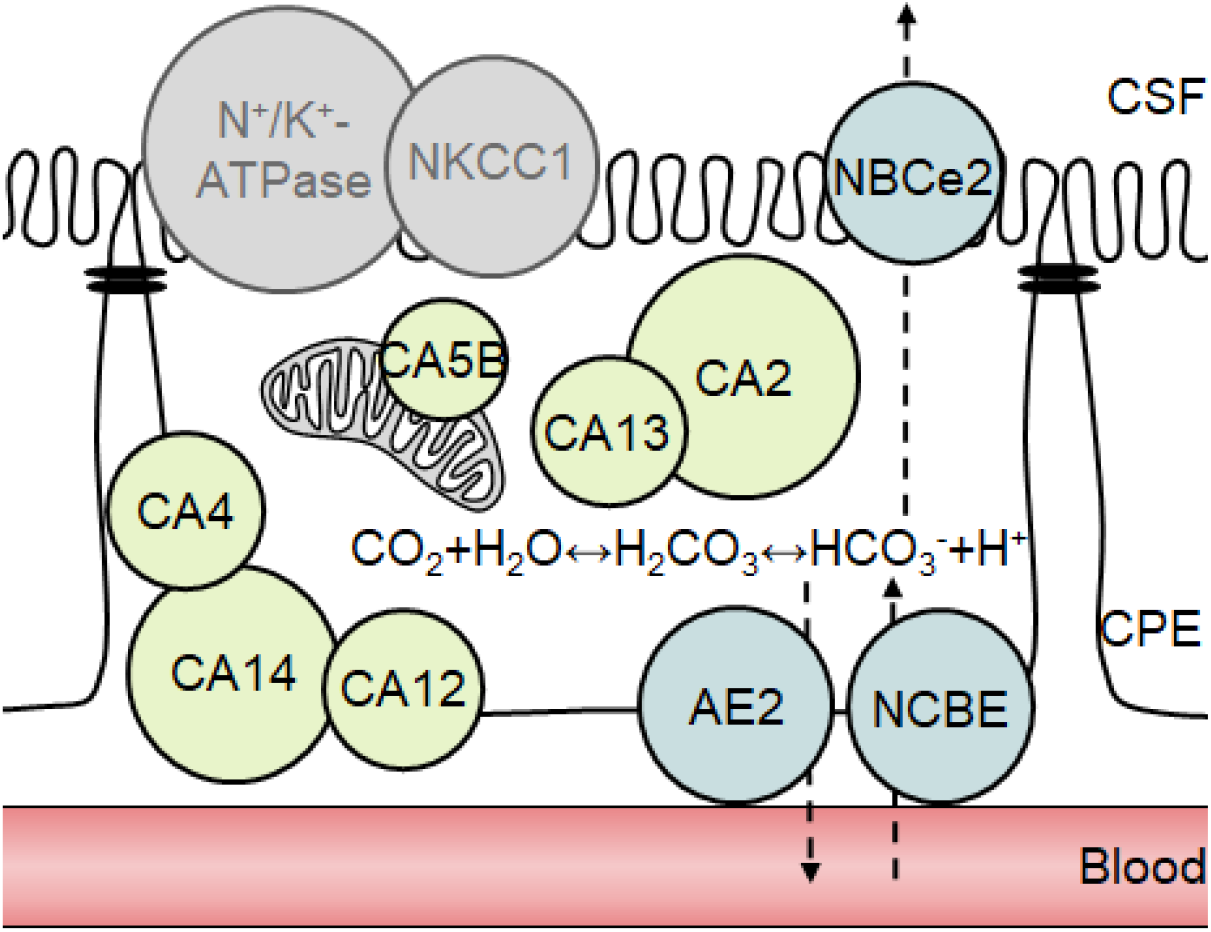
Schematic representation of carbonic anhydrases and transporters in a choroid plexus epithelial (CPE) cell. CA12 is expressed on the basolateral side of the choroid plexus (42), CA4, CA9 and CA14 are generally membrane bound (4), yet their exact localization (luminal or basolateral) in the choroid plexus cell remains unresolved. CA2 and CA13 are both cytosolic (4, 45), whereas CA5B is found in mitochondria (75). Not shown are CA6 and CA9, which are absent from mouse and human choroid plexus (Table 1), CA3, which is AZE insensitive (44), and CA11, which may be acatalytic (4). CA isoforms 1, 7, 8, 10 and 15 were bellow detection level in rat choroid plexus (Table 1). The key transporters involved in CSF secretion NKCC1 and Na/K-ATPase are not affected by AZE treatment. The main candidates for mediating AZE-induced reduction in CSF secretion are the bicarbonate transporters: AE2 and NBCE/NBCn2 expressed at the basolateral membrane and NBCe2 expressed at the luminal membrane. The sphere area indicates transcriptional expression levels.

Administration of AZE by the i.v. route caused an initial peak in ICP prior to the subsequent gradual decline. This peak was mirrored by a decrease in the exhaled CO_2_ and thus an elevated blood pCO_2_. Such abrupt increase in pCO_2_ may cause intracranial vasodilation (74), which could have caused the observed peak in ICP, similar to that observed upon a switch from 100% O_2_ inhalation to 30% CO_2_ in experimental cats (27). In support of its vascular origin, the elevated pCO_2_ and the resulting ICP peak was absent in the experimental rats that had AZE administered through the i.c.v. route. The AZE-mediated elevation in blood pCO_2_ upon systemic application remained throughout the duration of the experiment. Organisms usually hyperventilate to correct for pCO_2_ elevation. Mechanical hyperventilation of the experimental rats reduced the blood pCO_2_ in both control rats and those exposed to AZE compared to rats with ‘normal’ ventilation parameters. Nevertheless, the AZE-induced ICP reduction remained intact – it was even slightly more pronounced. The latter finding suggests that AZE treatment in combination with the hyperventilation, sometimes employed clinically to treat elevated ICP (54), may serve as complementary tools to manage ICP. However, this additive effect is possibly caused by secondary mechanisms, like pCO_2_-mediated reduction in cerebral blood flow (23), as it was shown that CSF secretion did not differ between animals with ‘normal’ ventilation and with hyperventilation (29). Taken together, AZE-mediated ICP reduction does not arise from the increase in pCO_2_.

In conclusion, AZE reduces the ICP in healthy rats via its ability to decrease the CSF secretion rate. AZE exerts its effect on the CSF secretion machinery in a direct manner, most likely by targeting the choroidal carbonic anhydrases. These enzymes modulate the substrate availability for the HCO_3_ transporters that are highly expressed in the choroid plexus and known to act as key contributors to CSF secretion across this tissue (35, 40, 41). A non-selective CA inhibitor like AZE affects carbonic anhydrases in all other tissues and cell types in the body, which causes the many unpleasant side effects observed with usage of this inhibitor (10). However, these actions do not, as such, appear to affect the CSF secretion rate or the ICP. These findings provide promise of future selective targeting of choroidal carbonic anhydrases in the search for a pharmacological approach to reduce ICP elevation in patients experiencing any of the many pathologies demonstrating this feature.

## Author contributions

N.M., D.B., E.K.O., C.R.R. designed the research; D.B., E.K.O., J.H.W., T.L.T.B., E.C., S.N.A., N.J.G. executed the experiments/analyzed data. N.M., D.B. drafted the manuscript. All authors participated in the finalization of the manuscript.

## Acknowledgements

We thank laboratory manager Trine Lind Devantier, Department of Neuroscience, Faculty of Health and Medical Sciences, University of Copenhagen for technical assistance. We also thank veterinarians Maria Mathilde Haugaard and Karsten Pharao Hammelev from the Department of Experimental Medicine, Faculty of Health and Medical Sciences, University of Copenhagen for their input on optimizing the survival surgical procedures.

