## Supplementary files for "Acetazolamide modulates intracranial pressure directly by its action on the cerebrospinal fluid secretion apparatus"

| | CONTROL | | | | | | | | | ACETAZOLAMIDE | | | | | | | | | 2way ANOVA p-value | |
|---|---|---|---|---|---|---|---|---|---|---|---|---|---|---|---|---|---|---|---|---|
| | Baseline | | | 1h | | | 2h | | | Baseline | | | 1h | | | 2h | | | | |
| | Average | SEM | n | Average | SEM | n | Average | SEM | n | Average | SEM | n | Average | SEM | n | Average | SEM | n | Time | Treatment |
| pH | 7.393 | 0.009 | 5 | 7.359 | 0.009 | 5 | 7.337 | 0.014 | 5 | 7.379 | 0.011 | 5 | 7.156 | 0.012 | 5 | 7.126 | 0.009 | 5 | <.001 | <.001 |
| $pCO_2$ | 5.26 | 0.14 | 5 | 5.15 | 0.12 | 5 | 5.38 | 0.13 | 5 | 5.21 | 0.16 | 5 | 9.40 | 0.15 | 5 | 10.05 | 0.18 | 5 | <.001 | <.001 |
| $pO_2$ | 13.8 | 0.5 | 5 | 13.8 | 0.6 | 5 | 14.0 | 0.6 | 5 | 14.4 | 1.3 | 5 | 15.5 | 0.8 | 5 | 14.5 | 0.6 | 5 | 0.215 | 0.402 |
| ctHb | 9.5 | 0.1 | 5 | 9.2 | 0.2 | 5 | 9.6 | 0.2 | 5 | 9.3 | 0.2 | 5 | 9.2 | 0.2 | 5 | 9.5 | 0.3 | 5 | 0.033 | 0.729 |
| $sO_2$ | 94.2 | 0.8 | 5 | 94.7 | 1.7 | 5 | 93.3 | 1.0 | 5 | 94.3 | 1.3 | 4 | 90.2 | 1.4 | 5 | 88.6 | 1.4 | 5 | <.001 | 0.151 |
| $FO_2Hb$ | 94.5 | 0.8 | 5 | 94.1 | 1.2 | 5 | 93.3 | 1.1 | 5 | 95.3 | 1.8 | 5 | 90.1 | 1.5 | 5 | 88.5 | 1.4 | 5 | <.001 | 0.181 |
| $K^+$ | 5.2 | 0.1 | 5 | 4.9 | 0.1 | 5 | 5.0 | 0.1 | 5 | 5.4 | 0.2 | 5 | 5.1 | 0.2 | 5 | 5.9 | 0.3 | 5 | 0.038 | 0.043 |
| $Na^+$ | 139 | 0.6 | 5 | 142 | 0.7 | 5 | 144 | 1.1 | 5 | 138 | 0.7 | 5 | 143 | 1.0 | 5 | 144 | 1.3 | 5 | <.001 | 0.77 |
| $Ca^{2+}$ | 0.87 | 0.02 | 5 | 0.84 | 0.01 | 5 | 0.85 | 0.02 | 5 | 0.79 | 0.08 | 5 | 0.78 | 0.06 | 5 | 0.82 | 0.04 | 5 | 0.28 | 0.389 |
| $Cl^-$ | 102 | 0.7 | 5 | 107 | 0.9 | 5 | 110 | 0.5 | 5 | 101 | 0.9 | 5 | 105 | 0.7 | 5 | 106 | 0.7 | 5 | <.001 | 0.047 |
| cGlu | 22.6 | 0.8 | 5 | 18.4 | 1.2 | 5 | 11.9 | 1.0 | 5 | 24.5 | 0.6 | 5 | 19.5 | 0.8 | 5 | 14.4 | 0.9 | 5 | <.001 | 0.09 |
| cLac | 1.2 | 0.2 | 5 | 1.6 | 0.2 | 5 | 2.0 | 0.2 | 5 | 1.1 | 0.2 | 5 | 0.8 | 0.1 | 5 | 0.9 | 0.1 | 5 | 0.223 | <.001 |
| ctBil | 53 | 2.6 | 5 | 48 | 1.8 | 4 | 54 | 3.8 | 5 | 50 | 1.2 | 3 | 39 | 1.2 | 3 | 42 | 1.4 | 3 | 0.001 | 0.09 |
| cBase | -0.9 | 0.5 | 5 | -3.7 | 0.5 | 5 | -4.2 | 0.7 | 5 | -2.2 | 0.6 | 5 | -3.9 | 0.6 | 5 | -4.4 | 0.5 | 5 | <.001 | 0.482 |
| $cHCO_3^-$ | 23.7 | 0.4 | 5 | 21.6 | 0.4 | 5 | 21.1 | 0.6 | 5 | 23.0 | 0.4 | 4 | 19.6 | 0.5 | 4 | 19.3 | 0.3 | 4 | <.001 | 0.04 |

**Supplementary Table S1** Blood gas analysis in intact animals before (baseline) and after (at 1 and 2 h) i.v. injection of 100 mg $kg^{-1}$ AZE in anesthetized and ventilated rats.

| | CONTROL NEPHRECTOMY | | | | | | | | | ACETAZOLAMIDE NEPHRECTOMY | | | | | | | | | 2way ANOVA p-value | |
|---|---|---|---|---|---|---|---|---|---|---|---|---|---|---|---|---|---|---|---|---|
| | Baseline | | | 1h | | | 2h | | | Baseline | | | 1h | | | 2h | | | | |
| | Average | SEM | n | Average | SEM | n | Average | SEM | n | Average | SEM | n | Average | SEM | n | Average | SEM | n | Time | Treatment |
| pH | 7.343 | 0.015 | 4 | 7.295 | 0.020 | 4 | 7.264 | 0.012 | 4 | 7.368 | 0.018 | 4 | 7.148 | 0.013 | 4 | 7.100 | 0.018 | 4 | <.001 | 0.005 |
| $pCO_2$ | 5.34 | 0.17 | 4 | 4.78 | 0.26 | 4 | 5.41 | 0.09 | 4 | 4.99 | 0.08 | 4 | 9.08 | 0.21 | 4 | 10.16 | 0.29 | 4 | <.001 | <.001 |
| $pO_2$ | 16.4 | 0.6 | 4 | 16.4 | 0.7 | 4 | 14.5 | 1.7 | 4 | 16.3 | 1.5 | 4 | 17.1 | 0.7 | 4 | 16.5 | 1.0 | 4 | 0.303 | 0.539 |
| ctHb | 9.4 | 0.1 | 3 | 9.0 | 0.2 | 4 | 9.0 | 0.1 | 4 | 9.6 | 0.5 | 4 | 9.2 | 0.4 | 4 | 8.7 | 0.2 | 4 | 0.136 | 0.761 |
| $sO_2$ | 97.2 | 1.9 | 3 | 96.3 | 1.8 | 4 | 93.0 | 3.4 | 4 | 96.4 | 2.0 | 4 | 94.5 | 1.3 | 4 | 91.2 | 1.3 | 4 | 0.033 | 0.719 |
| $FO_2Hb$ | 96.9 | 1.5 | 3 | 95.6 | 1.1 | 4 | 92.2 | 3.0 | 4 | 95.7 | 1.4 | 4 | 94.8 | 1.6 | 4 | 91.9 | 0.9 | 3 | 0.059 | 0.819 |
| $K^+$ | 5.8 | 0.4 | 4 | 5.1 | 0.1 | 4 | 6.0 | 0.2 | 4 | 5.6 | 0.0 | 4 | 5.7 | 0.2 | 4 | 6.2 | 0.2 | 4 | 0.025 | 0.382 |
| $Na^+$ | 138 | 0.8 | 4 | 143 | 0.6 | 4 | 144 | 0.4 | 4 | 139 | 1.1 | 4 | 143 | 0.3 | 4 | 145 | 0.4 | 4 | <.001 | 0.072 |
| $Ca^{2+}$ | 0.84 | 0.05 | 4 | 0.82 | 0.04 | 4 | 0.85 | 0.06 | 4 | 0.84 | 0.02 | 4 | 0.71 | 0.05 | 4 | 0.77 | 0.05 | 4 | 0.033 | 0.321 |
| $Cl^-$ | 102 | 1.8 | 4 | 108 | 0.9 | 4 | 109 | 0.8 | 4 | 102 | 0.5 | 4 | 105 | 0.9 | 4 | 106 | 1.0 | 4 | <.001 | 0.062 |
| cGlu | 24.3 | 2.8 | 4 | 14.7 | 1.6 | 4 | 11.0 | 1.3 | 4 | 24.8 | 1.6 | 4 | 20.4 | 1.6 | 4 | 14.8 | 0.9 | 4 | <.001 | 0.132 |
| cLac | 1.4 | 0.2 | 4 | 1.9 | 0.1 | 4 | 1.9 | 0.2 | 4 | 1.0 | 0.1 | 4 | 0.7 | 0.1 | 4 | 0.8 | 0.1 | 4 | 0.344 | 0.003 |
| ctBil | 54 | 1.4 | 2 | 45 | 0.3 | 3 | 41 | 2.9 | 3 | 56 | 5.7 | 3 | 42 | 3.2 | 2 | 40 | 1.1 | 2 | | |
| cBase | -4.0 | 0.6 | 4 | -9.2 | 0.6 | 4 | -8.7 | 0.7 | 4 | -3.8 | 1.1 | 4 | -5.3 | 0.4 | 4 | -6.1 | 0.7 | 4 | <.001 | 0.043 |
| $cHCO_3^-$ | 21.4 | 0.5 | 4 | 17.8 | 0.5 | 4 | 17.8 | 0.5 | 4 | 21.7 | 0.8 | 4 | 19.0 | 0.4 | 4 | 18.2 | 0.5 | 4 | <.001 | 0.424 |

**Supplementary Table S2** Blood gas analysis before (baseline) and after (at 1 and 2 h) i.v. injection of 100 mg $kg^{-1}$ AZE in in anesthetized, ventilated and nephrectomized rats

| | CONTROL HYPERVENTILATION | | | | | | | | | ACETAZOLAMIDE HYPERVENTILATION | | | | | | | | | 2way ANOVA p-value | |
|---|---|---|---|---|---|---|---|---|---|---|---|---|---|---|---|---|---|---|---|---|
| | Baseline | | | 1h | | | 2h | | | Baseline | | | 1h | | | 2h | | | | |
| | Average | SEM | n | Average | SEM | n | Average | SEM | n | Average | SEM | n | Average | SEM | n | Average | SEM | n | Time | Treatment |
| pH | 7.434 | 0.010 | 4 | 7.585 | 0.025 | 4 | 7.525 | 0.041 | 4 | 7.425 | 0.012 | 4 | 7.216 | 0.018 | 4 | 7.150 | 0.025 | 4 | 0.001 | <.001 |
| $pCO_2$ | 4.77 | 0.11 | 4 | 2.04 | 0.06 | 4 | 1.92 | 0.05 | 4 | 4.84 | 0.10 | 4 | 7.80 | 0.15 | 4 | 8.37 | 0.32 | 4 | 0.039 | <.001 |
| $pO_2$ | 14.4 | 1.2 | 4 | 22.8 | 0.3 | 4 | 22.8 | 0.3 | 4 | 13.8 | 0.6 | 4 | 23.0 | 0.2 | 4 | 23.1 | 0.4 | 4 | <.001 | 0.931 |
| ctHb | 9.3 | 0.1 | 4 | 9.3 | 0.3 | 4 | 9.8 | 0.3 | 4 | 9.8 | 0.2 | 4 | 9.7 | 0.2 | 4 | 10.4 | 0.3 | 4 | 0.022 | 0.128 |
| $sO_2$ | 95.3 | 0.7 | 4 | NA | | | NA | | | 95.6 | 1.6 | 4 | 97.7 | 1.2 | 4 | 97.3 | 1.3 | 4 | | |
| $FO_2Hb$ | 95.5 | 0.7 | 4 | NA | | | NA | | | 95.0 | 1.0 | 4 | 97.4 | 0.6 | 4 | 97.2 | 0.8 | 4 | | |
| $K^+$ | 5.2 | 0.1 | 4 | 5.2 | 0.1 | 4 | 4.9 | 0.1 | 4 | 5.4 | 0.2 | 4 | 5.8 | 0.2 | 4 | 6.2 | 0.2 | 4 | 0.185 | 0.009 |
| $Na^+$ | 139 | 0.9 | 4 | 140 | 0.5 | 4 | 142 | 1.7 | 4 | 139 | 0.0 | 4 | 142 | 0.4 | 4 | 143 | 0.8 | 4 | 0.007 | 0.326 |
| $Ca^{2+}$ | 0.92 | 0.02 | 4 | 0.88 | 0.01 | 4 | 0.86 | 0.03 | 4 | 0.87 | 0.02 | 4 | 0.86 | 0.01 | 4 | 0.82 | 0.02 | 4 | 0.094 | 0.056 |
| $Cl^-$ | 102 | 1.1 | 4 | 106 | 1.1 | 4 | 110 | 1.6 | 4 | 102 | 0.5 | 4 | 105 | 0.9 | 4 | 108 | 0.6 | 4 | <.001 | 0.377 |
| cGlu | 19.5 | 1.5 | 4 | 21.7 | 1.6 | 4 | 16.0 | 1.9 | 4 | 22.7 | 1.1 | 4 | 21.9 | 1.9 | 4 | 19.5 | 2.2 | 4 | 0.001 | 0.352 |
| cLac | 1.1 | 0.1 | 4 | 4.2 | 0.5 | 4 | 4.6 | 0.4 | 4 | 1.1 | 0.1 | 4 | 0.8 | 0.1 | 4 | 1.0 | 0.1 | 4 | <.001 | <.001 |
| ctBil | 51 | 1.6 | 4 | NA | | | NA | | | 53 | 3.6 | 3 | 53 | 4.2 | 2 | 63 | 2.8 | 2 | | |
| cBase | -0.3 | 1.2 | 4 | -7.3 | 1.3 | 4 | -10.8 | 1.8 | 4 | -0.7 | 0.6 | 4 | -4.1 | 0.9 | 4 | -7.1 | 0.9 | 4 | <.001 | 0.205 |
| $cHCO_3^-$ | 24.5 | 0.9 | 4 | NA | | | NA | | | 24.1 | 0.5 | 4 | 20.3 | 0.7 | 4 | 17.9 | 0.7 | 4 | | |

**Supplementary Table S3** Blood gas analysis before (baseline) and after (at 1 and 2 h) i.v. injection of 100 mg $kg^{-1}$ AZE in anesthetized and hyperventilated rats. NA – the values were over the detection threshold.

| | CONTROL INTRAVENTRICULAR | | | | | | | | | ACETAZOLAMIDE INTRAVENTRICULAR | | | | | | | | | 2way ANOVA p-value | |
|---|---|---|---|---|---|---|---|---|---|---|---|---|---|---|---|---|---|---|---|---|
| | Baseline | | | 1h | | | 2h | | | Baseline | | | 1h | | | 2h | | | | |
| | Average | SEM | n | Average | SEM | n | Average | SEM | n | Average | SEM | n | Average | SEM | n | Average | SEM | n | Time | Treatment |
| pH | 7.409 | 0.014 | 4 | 7.367 | 0.017 | 4 | 7.357 | 0.021 | 4 | 7.421 | 0.010 | 4 | 7.387 | 0.002 | 4 | 7.358 | 0.007 | 4 | <.001 | 0.526 |
| $pCO_2$ | 4.76 | 0.18 | 4 | 4.47 | 0.12 | 4 | 4.76 | 0.14 | 4 | 4.77 | 0.08 | 4 | 4.70 | 0.15 | 4 | 4.78 | 0.10 | 4 | 0.230 | 0.551 |
| $pO_2$ | 16.3 | 1.7 | 4 | 15.8 | 0.7 | 4 | 14.9 | 0.7 | 4 | 15.2 | 1.4 | 4 | 14.9 | 0.9 | 4 | 15.0 | 0.9 | 4 | 0.388 | 0.649 |
| ctHb | 9.3 | 0.3 | 4 | 9.5 | 0.2 | 4 | 9.6 | 0.2 | 4 | 9.5 | 0.3 | 4 | 9.9 | 0.2 | 4 | 9.9 | 0.1 | 4 | 0.167 | 0.289 |
| $sO_2$ | 95.6 | 0.7 | 4 | 95.2 | 0.5 | 4 | 94.4 | 1.1 | 4 | 95.0 | 1.0 | 4 | 94.0 | 0.8 | 4 | 93.9 | 0.7 | 4 | 0.089 | 0.487 |
| $FO_2Hb$ | 96.0 | 0.7 | 4 | 95.3 | 0.6 | 4 | 94.6 | 1.1 | 4 | 95.1 | 1.0 | 4 | 94.2 | 0.8 | 4 | 94.1 | 0.7 | 4 | 0.074 | 0.487 |
| $K^+$ | 5.2 | 0.2 | 4 | 4.8 | 0.3 | 4 | 5.0 | 0.2 | 4 | 5.2 | 0.1 | 4 | 5.0 | 0.2 | 4 | 5.2 | 0.3 | 4 | 0.263 | 0.582 |
| $Na^+$ | 140 | 0.3 | 4 | 144 | 1.0 | 4 | 144 | 0.4 | 4 | 141 | 1.0 | 4 | 143 | 1.1 | 4 | 145 | 0.0 | 4 | 0.007 | 0.55 |
| $Ca^{2+}$ | 0.88 | 0.05 | 4 | 0.85 | 0.04 | 4 | 0.85 | 0.03 | 4 | 0.89 | 0.03 | 4 | 0.86 | 0.03 | 4 | 0.86 | 0.03 | 4 | 0.081 | 0.895 |
| $Cl^-$ | 104 | 1.8 | 4 | 110 | 1.6 | 4 | 108 | 2.3 | 4 | 101 | 1.3 | 4 | 106 | 0.6 | 4 | 109 | 1.7 | 4 | 0.024 | 0.204 |
| cGlu | 19.8 | 1.3 | 4 | 13.1 | 0.6 | 4 | 8.4 | 0.8 | 4 | 21.4 | 1.7 | 4 | 17.5 | 1.1 | 4 | 9.9 | 0.4 | 4 | <.001 | 0.081 |
| cLac | 1.5 | 0.1 | 4 | 2.3 | 0.2 | 4 | 1.9 | 0.2 | 4 | 1.1 | 0.1 | 4 | 1.9 | 0.2 | 4 | 2.0 | 0.1 | 4 | 0.001 | 0.313 |
| ctBil | 51 | 2.5 | 4 | 51 | 2.7 | 4 | 55 | 1.9 | 4 | 52 | 3.0 | 4 | 57 | 1.8 | 4 | 59 | 1.8 | 4 | <.001 | 0.284 |
| cBase | -2.1 | 1.7 | 4 | -6.1 | 1.1 | 4 | -5.5 | 1.2 | 4 | -1.3 | 0.4 | 4 | -3.8 | 0.6 | 4 | -5.4 | 0.8 | 4 | 0.001 | 0.447 |
| $cHCO_3^-$ | 23.1 | 1.2 | 4 | 19.5 | 1.0 | 4 | 18.2 | 2.1 | 4 | 23.7 | 0.3 | 4 | 21.8 | 0.3 | 4 | 20.6 | 0.6 | 4 | 0.001 | 0.265 |

**Supplementary Table S4** Blood gas analysis before (baseline) and after (at 1 and 2 h) the i.c.v. infusion of 18 mM AZE (expected ventricular concentration 500 µM) in anesthetized and ventilated rats.

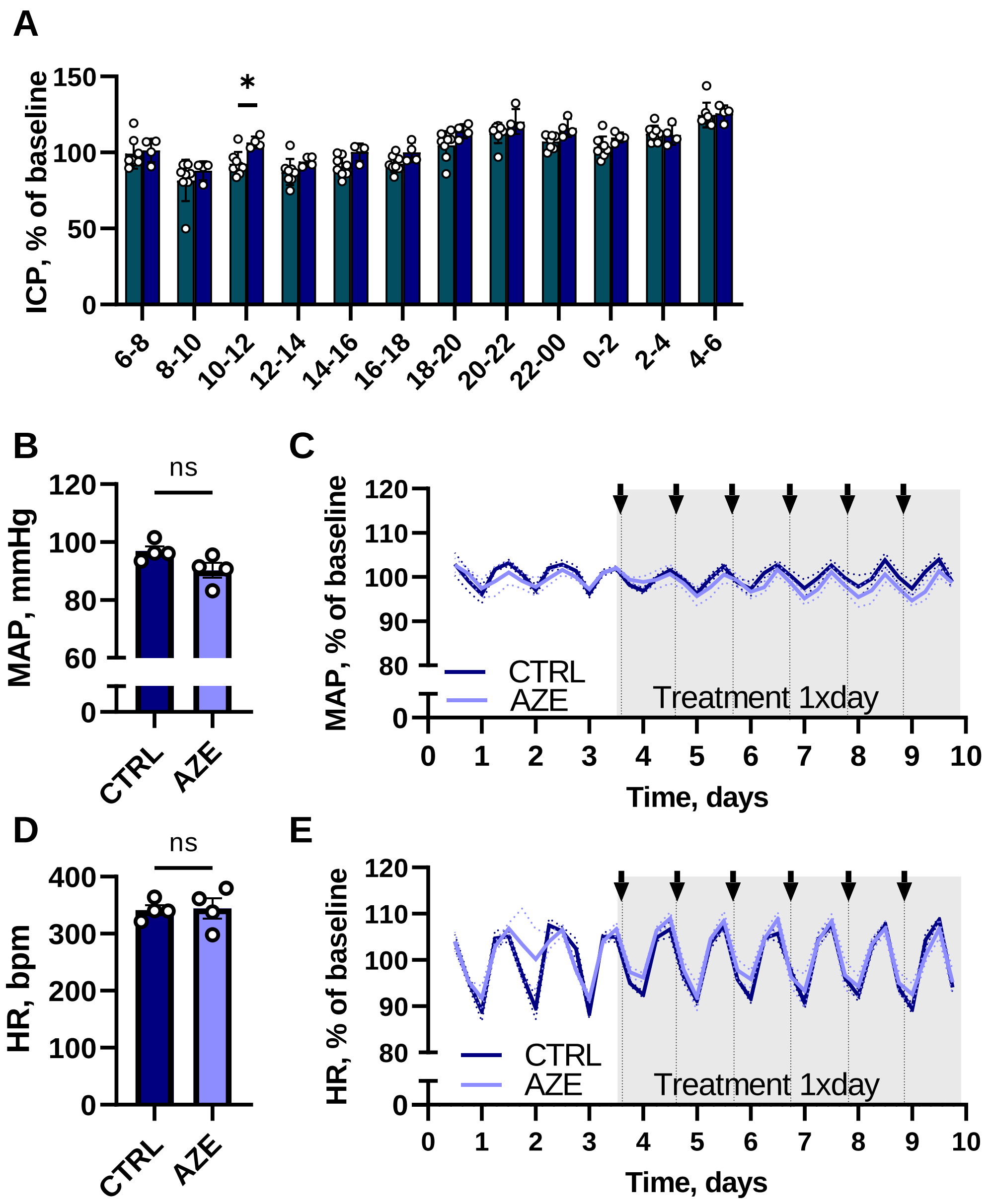


**Supplementary Figure 1 Telemetric measurements in awake rats.** Comparison between the baseline daily ICP fluctuations and p.o. saline treatement is shown in **A**, by presenting the daily 2 h average ICP normalized to the 24h baseline value during the baseline period (n = 8), and during once daily treatment with p.o. saline (n = 4). 2way ANOVA with Bonferroni's multiple comparisons test showed significant difference only during the first 2h after administration. Figure **B** represents the 72 h average MAP, separated between the animals that afterwards received either 1 × daily 100 mg $kg^{-1}$ p.o. AZE treatment or control ($MAP_{CTRL}$ = 97 ± 2 mmHg, $MAP_{AZE}$ = 90 ± 3 mmHg, n = 4 in each group, P = 0.07). The daily MAP fluctuations, normalized to the 24h baseline MAP are shown as 4h bins in **C**. The heart rate data is shown in **D** ($HR_{CTRL}$ = 342 ± 9 bpm, $HR_{AZE}$ = 344 ± 18 bpm, n = 4 in each group, P = 0.9) and **E** in the same manner as the MAP.
